# Viral satellites exploit phage proteins to escape degradation of the bacterial host chromosome

**DOI:** 10.1101/671271

**Authors:** Amelia C. McKitterick, Stephanie G. Hays, Munirul Alam, Kimberley D. Seed

## Abstract

Phage defense systems are often found on mobile genetic elements (MGEs), where they constitutively defend against invaders or are induced to respond to new assaults. Some MGEs, the phage satellites, exploit phages for their own transmission after induction, reducing phage production and protecting their hosts in the process. One such satellite in *Vibrio cholerae*, PLE, is triggered by the lytic phage ICP1 to excise from the chromosome, replicate, and transduce to neighboring cells, completely sabotaging phage production. Here, we found that ICP1 has evolved to possess one of two syntenic loci encoding an SF1B-type helicase, either of which PLE can exploit to directly drive PLE replication. Further, loss of PLE mobilization limits anti-phage activity due to phage-mediated degradation of the bacterial genome. Our work provides insight into the unique challenges imposed on the parasites of lytic phages and underscores the adaptions of these satellites to their ever-evolving target phage.

## Introduction

Viruses and mobile genetic elements (MGEs) are associated with organisms from all branches of the tree of life (Koonin & Krupovic 2015). In order to successfully infect their hosts, viruses employ a variety of host-takeover programs that inhibit host activities while promoting viral processes. Bacteriophages, or phages, are viruses that infect bacterial hosts and have profound effects on bacterial fitness, as well as on human health and disease (Brüssow et al. 2004; Bondy-Denomy & Davidson 2014). Of interest, lytic phages, which infect and kill their bacterial hosts within a single round of infection, have recently come to light as having impactful roles in shaping the composition of bacterial populations, such as the human gut microbiome (Manrique et al. 2017), and as potential biocontrol agents for antibiotic resistant infections (Pires et al. 2016). Lytic phages are particularly insidious to their bacterial hosts—upon infection, phages like the lytic *Escherichia coli* phage T4 can express a variety of genes that mediate host-cell takeover programs. T4 expresses genes that shut down and redirect host transcriptional machinery to favor transcription of phage genes, as well as nucleases that degrade the host chromosome to inhibit host gene expression as well as free up nucelosides that are then incorporated into the rapidly replicating phage genome (Hinton et al. 2005; Warner et al. 1970; Hercules et al. 1971).

Paradoxically, phages also contribute to bacterial population diversity and complexity by facilitating horizontal gene transfer (HGT) (Brüssow et al. 2004; Koskella & Brockhurst 2014). In addition to the well characterized mechanisms by which phages can spread bacterial genetic material to neighboring cells, such as generalized and specialized transduction (Penadés et al. 2015), recent work has uncovered a means for large regions of the bacterial chromosome to be packaged into temperate phage virions in a process termed lateral transduction (Chen et al. 2018). Independent of packaging, phages can also facilitate the spread of bacterial plasmid DNA from lysed cells to neighbors, increasing the range of genetic material that can be shared within a population (Keen et al. 2017). In sharp contrast to these forms of “passive” phage-mediated HGT, there are parasitic mobile genetic elements referred to as phage satellites, such as <u>p</u>hage <u>i</u>nducible <u>c</u>hromosomal <u>i</u>slands (PICIs), that have evolved to explicitly manipulate the phage replication and packaging programs for their own horizontal spread (Penadés & Christie 2015).

Typically, phage defense in bacteria is attributable to widely characterized systems, including restriction-modification and CRISPR-Cas systems that inhibit phage through targeted cleavage of the infecting phage genome, and toxin/antitoxin systems and abortive infection systems that function through killing of the infected host cell (Samson et al. 2013; Dy et al. 2014; Hille et al. 2018). However, phage parasites, which are being increasingly discovered (O’Hara et al. 2017; Martínez-Rubio et al. 2017; Fillol-Salom et al. 2018), can also provide robust phage defense for their bacterial hosts. One type of PICI, the well characterized *<u>S</u>taphlococcus <u>a</u>ureus* <u>p</u>athogenicity <u>i</u>slands (SaPIs), are induced by infection with a helper phage, compete with that helper over the bacterial host’s replication machinery, and steal phage packaging proteins to selfishly package the SaPI genome for horizontal transfer. This parasitic interference negatively impacts the ability of the helper phage to complete its lifecycle, thus blocking plaque formation (Ubeda et al. 2009; Tormo-Más et al. 2010; Ram et al. 2012). Despite diverse mechanisms, phage defense systems must overcome phage-mediated host takeover and go on to prevent rampant phage propagation through the bacterial community. Genomic analyses to localize anti-phage mechanisms in bacterial genomes have revealed that they tend to cluster together on what are referred to as defense islands (DIs) (Makarova et al. 2011). Analysis of DIs has even led to the discovery of new phage defense systems solely due to the prevalent clustering of these defense systems on MGEs (Doron et al. 2018). While hypothesized to have roles in HGT, the prevalence of phage defense systems on genomic islands has yet to be explained. Likewise, it remains to be seen the extent to which such DIs have evolved to parasitize phages for their own dissemination.

*Vibrio cholerae*, the etiological agent of the diarrheal disease cholera, is constantly under assault by phages both in aquatic environments as well as in human hosts (Faruque et al. 2005; Seed et al. 2011; Seed et al. 2014). The dominant phage that preys on epidemic *V. cholerae* is ICP1, a lytic myovirus that is consistently isolated from cholera patient stool samples in regions where cholera is endemic, such as Dhaka, Bangladesh (Seed et al. 2011; Angermeyer et al. 2018; McKitterick et al. 2019). In response to the consistent attack by ICP1, *V. cholerae* has acquired the <u>p</u>hage-inducible chromosomal island-<u>l</u>ike <u>e</u>lement (PLE), a highly specific phage satellite that blocks plaque formation by ICP1 while exploiting phage resources to further its lifecycle (O’Hara et al. 2017). PLE excises from the host chromosome during ICP1 infection, replicates to high copy, and is specifically transduced to neighboring cells; concurrently, PLE replication within the host cell negatively impacts the ability of ICP1 to replicate its genome, contributing to the inhibition of ICP1 production (Barth et al. 2019). PLE encodes a large serine recombinase, Int, that catalyzes the PLE excision and circularization reaction by physically interacting with ICP1-encoded PexA, a small protein of unknown function that is specific to ICP1 and is hijacked by PLE to act as a recombination directionality factor (McKitterick & Seed 2018). Once excised, PLE begins to replicate and then is thought to steal structural proteins from ICP1 to facilitate its own transmission. Once packaged, PLE triggers accelerated lysis of the infected culture allowing for release of PLE transducing particles from the infected cells, ultimately killing the infected *V. cholerae* host but protecting the population as no infectious ICP1 progeny are produced (O’Hara et al. 2017). Five PLEs have been identified in epidemic *V. cholerae* isolates and all block plaque formation by ICP1. In addition to inhibiting ICP1, PLEs are also characterized by conserved genomic architecture and the aforementioned PLE lifecycle during ICP1 infection (O’Hara et al. 2017).

Recent work has uncovered a PLE-encoded factor that is necessary for PLE replication: the replication initiation factor, *repA* (Barth et al. 2019). Expression of *repA* is induced by ICP1 infection (unpublished), facilitating origin binding and recruitment of replisome proteins that have yet to be identified. In the absence of ICP1 infection, however, RepA is not sufficient to drive PLE replication (Barth et al. 2019), further, PLE is not predicted to encode replication machinery, suggesting that other phage-encoded gene products are required for PLE amplification. As all PLEs replicate following ICP1 infection (O’Hara et al. 2017), it stands to reason that the PLE has evolved to exploit conserved components of ICP1’s replication machinery. Similar to PLE excision (O’Hara et al. 2017), PLE replication is essential for horizontal transmission of PLE transducing units, thus further underscoring the role of ICP1 in driving PLE HGT; however, the relatively low rate of transduction suggests that robust PLE replication may have other roles in the PLE conflict with ICP1 (Barth et al. 2019).

In order to exploit ICP1, PLE must escape from ICP1-mediated host takeover during infection. While the precise mechanisms that ICP1 uses to overcome *V. cholerae* have not been characterized, ICP1 is able to rapidly begin replicating its genome following infection (Barth et al. 2019) and produces about 100 virions within 20 minutes of infection (O’Hara et al. 2017). Here, we identify ICP1 *ΔpexA* mutants that escape PLE by acquiring mutations in the ICP1-encoded SF1B accessory helicase that we have named *helA*. We show that while this helicase is not necessary for ICP1 replication, it is essential for PLE to hijack to drive its own replication during ICP1 infection. We show that the excision- and replication-deficient PLE is susceptible to ICP1-mediated host takeover, whereby PLE is degraded while it remains integrated in the *V. cholerae* chromosome. Analysis of natural isolates of ICP1 from cholera patient stool samples in the megacity of Dhaka compared to a rural site in Bangladesh revealed an alternative SF1B helicase allele in phages shed from the rural site. Functional comparisons between the two alleles revealed that both alleles, though unrelated, can be hijacked by all PLEs to facilitate PLE replication. Though neither helicase is essential for ICP1, ICP1 faces impaired fitness in the absence of either accessory helicase, explaining their prevalence in ICP1 and other *Vibrio* phages. PLE’s capacity to use a variety of phage-encoded helicases to drive PLE replication underscores the critical role that replication plays in the PLE lifecycle to avoid phage-mediated host takeover and to facilitate continued gene expression. The common trend of phage defense islands clustering on MGEs suggests that mobilization of these phage defense islands, such as PLE, is a common mechanism to escape phage mediated host takeover.

## Results

### ICP1 is able to escape excision deficient PLE by acquiring mutations in the predicted helicase *helA*

Previous work has demonstrated the role for phage-encoded *pexA* in directing PLE 1 excision during infection with an ICP1 isolate from 2006, referred to as ICP1^A^ (McKitterick & Seed 2018) (Figure 1A). PLE 1 mediated inhibition of ICP1 does not require PLE 1 excision, so ICP1 *ΔpexA* is still blocked by PLE 1 (Figure 1B); however, ICP1^A^ *ΔpexA* is able to form rare plaques on *V. cholerae* harboring PLE 1 at a frequency of about 1 per 10^6^ phage (Figures 1A and 1C). Due to the low efficiency of plaquing, we consider these phage to be “escape phage” that have acquired a mutation in the genome allowing them to overcome PLE 1. To identify the phage gene(s) that harbor mutations enabling escape, we collected and purified three escape phage and performed whole genome sequencing. Analysis of these genomes revealed that all escape phage had acquired mutations in ICP1^A^ *gp147*, a predicted SF1B-type helicase which we have since named *<u>heli</u>case <u>A</u> (helA)* (Table S1).

**Figure 1.**
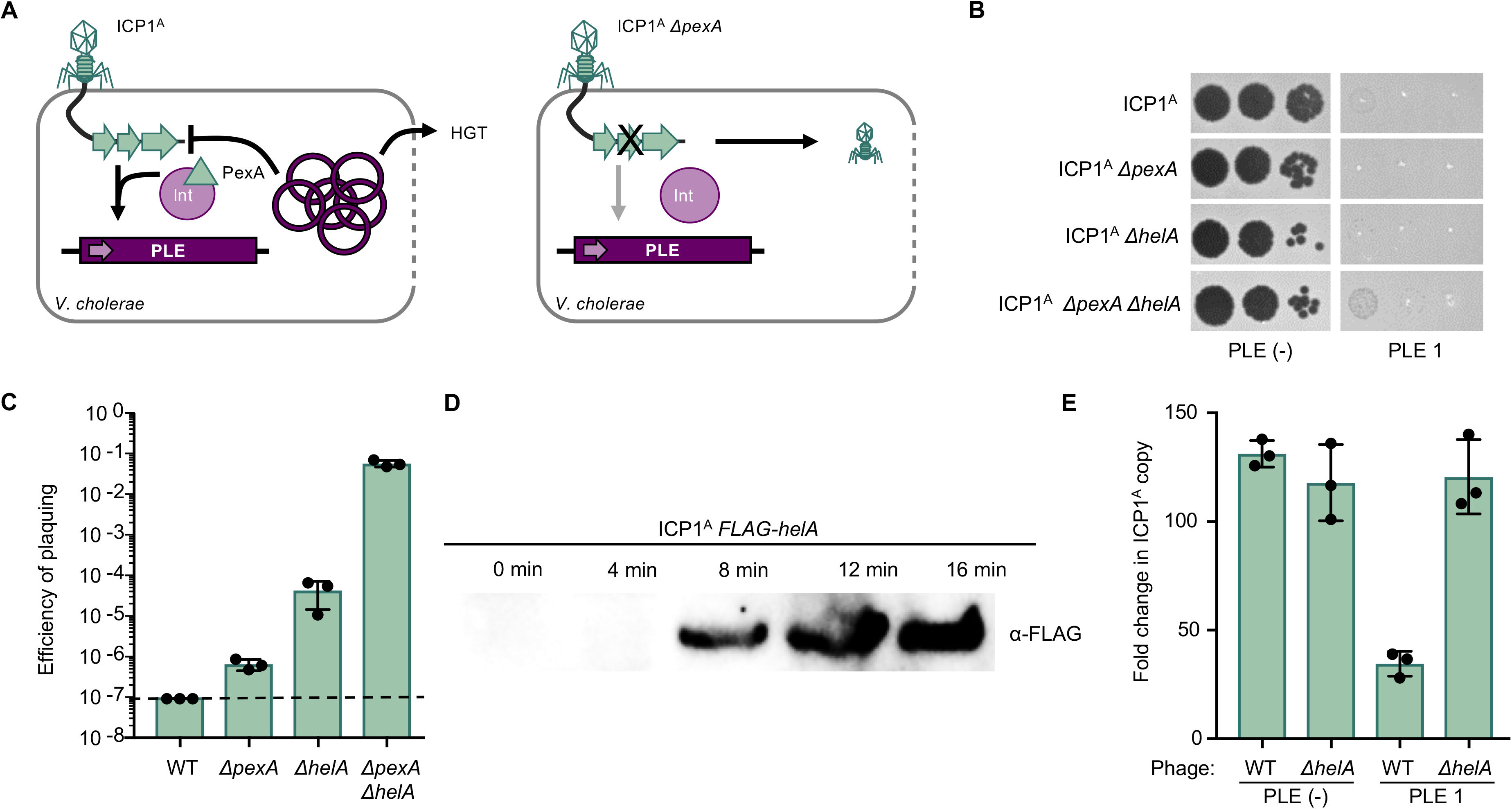
ICP1 overcomes excision-deficient PLE through loss of accessory helicase *helA*. **A**, Schematic of the PLE 1 response to ICP1 infection. Left, ICP1 infects PLE (+) *V. cholerae* and expresses PexA, which physically interacts with PLE 1-encoded integrase (Int) to direct PLE circularization and excision. Excised PLE 1 replicates to high copy number, inhibits ICP1 replication, and horizontally transduces to neighboring cells when *V. cholerae* undergoes PLE 1-mediated accelerated lysis. Right, when ICP1 *ΔpexA* infects PLE (+) *V. cholerae*, PLE 1 remains integrated in the host chromosome, and rare mutant phage are able to escape and form a plaque. **B**, Tenfold dilutions of ICP1 spotted on a PLE 1 and PLE (−) *V. cholerae* lawn (grey). Zones of killing are shown in black. **C**, Efficiency of plaquing of wild-type (WT) ICP1^A^ or derivatives with the deletions listed on PLE 1 relative to a PLE (−) *V. cholerae* host. Dashed line indicates limit of detection. **D**, Western blot of endogenously FLAG-tagged HelA during infection of PLE (−) *V. cholerae*. **E**, Quantification of change in ICP1 genome copy number following 20 minutes of infection of the listed *V. cholerae* host as detected by qPCR.

SF1B-type helicases are found broadly across all domains of life and include the well-studied *pif1* and *recD* (Saikrishnan et al. 2009). In eukaryotes such as *Saccharomyces cerevisiae, pif1* has been implicated in telomere maintenance, Okazaki fragment processing, and resolution of G-quadruplex motifs (Byrd & Raney 2017), while *recD* is a core component of the *E. coli* RecBCD complex involved in DNA processing and repair (Singleton et al. 2004). Another prototypical SF1B-type helicase is *dda*, encoded by phage T4, which is a non-essential accessory helicase implicated in origin melting, translocating proteins off DNA, and a wide variety of other functions *in vitro*, although its exact role *in vivo* is unknown (He et al. 2012; Byrd & Raney 2006; Brister 2008).

To validate the role of *helA* in ICP1^A^ escape from PLE 1, we constructed a *helA* deletion in a wild-type ICP1^A^ background and probed the mutant phage for the ability to overcome PLE 1. ICP1^A^-encoded *helA* is not necessary for plaque formation on PLE (−) *V. cholerae*, and ICP1^A^ *ΔhelA* is still blocked by PLE 1, indicating that *helA* is not necessary for PLE 1 induction (Figure 1B). Similar to ICP1^A^ *ΔpexA*, the absence of *helA* gives ICP1^A^ an advantage on PLE (+) *V. cholerae*, allowing for rare plaques to form; however, ICP1^A^*ΔhelA* forms plaques at a frequency two orders of magnitude higher than ICP1^A^ *ΔpexA* on PLE 1 (Figure 1C). Conversely, the double mutant ICP1^A^ *ΔpexA ΔhelA* is able to form small plaques on PLE 1 at a relatively high efficiency (Figures 1B and 1C). ICP1^A^ *ΔpexA ΔhelA* plaques that form on PLE (+) *V. cholerae* were picked and the plaquing efficiency was re-tested to determine if those phage were subsequently able to escape PLE 1 at a higher rate (Figure S1). As these progeny phage re-plaqued at the same efficiency as ICP1^A^ *ΔpexA ΔhelA*, we conclude that they are not genetic escape phage but instead are able to overcome some aspects of PLE 1 activity through the loss of both ICP1^A^-encoded *pexA* and *helA*.

We next wanted to characterize the role of *helA* for ICP1^A^ function. HelA is detectable in infected cells via Western blot within eight minutes of ICP1^A^ infection (Figure 1D), which is consistent with the onset of ICP1^A^ replication initiation (Barth et al. 2019), suggesting that *helA* may have a role in ICP1^A^ replication. As PLE 1 diminishes the level of ICP1^A^ replication (O’Hara et al. 2017; Barth et al. 2019), we hypothesized that PLE 1 hijacks HelA during infection as a mechanism to interfere with ICP1^A^ replication. To test this hypothesis, we evaluated ICP1^A^ *ΔhelA* replication in the presence and absence of PLE 1 by qPCR. In contrast to plaque formation, which requires multiple rounds of phage infection and replication to visualize a zone of killing, qPCR allows for quantification of phage DNA replication in a single round of infection. Consistent with the ability to form a plaque on PLE (−) *V. cholerae*, there are no deficiencies in ICP1^A^ *ΔhelA* replication relative to a wild-type phage over the course of the 20 minute infection cycle (Figure 1E), indicating that *helA* is not essential for ICP1^A^ replication. Conversely, infection of a PLE (+) *V. cholerae* host with ICP1^A^ *ΔhelA* rescues ICP1 replication to the level that is observed in a PLE (−) host (Figure 1E), suggesting that, while not necessary for ICP1^A^, *helA* is exploited by PLE 1 to interfere with ICP1 during infection. However, because ICP1^A^ *ΔhelA* is not deficient for replication in the absence PLE 1, the ICP1^A^ replication defect in the presence of PLE 1 is not likely directly due to PLE 1-mediated hijacking of HelA activity.

### ICP1-encoded *helA* is necessary for PLE replication

ICP1 and PLE 1 replication appear to be inversely related, wherein ICP1 copy number is restored when PLE 1 replication is abolished via deletion of either the PLE 1 origin of replication or *repA* (Barth et al. 2019). Therefore, the observed restoration in ICP1^A^ *ΔhelA* copy number during infection of a PLE (+) host implicates phage-encoded HelA in promoting PLE 1 replication. To test the role of *helA* in PLE replication, we infected PLE (+) *V. cholerae* with ICP1^A^ *ΔhelA* and monitored the change in PLE 1 copy over the course of infection. While PLE 1 is able to replicate to high copy when infected with a wild-type phage, strikingly, PLE 1 is unable to replicate in the absence of *helA* (Figure 2A). This phenotype can be complemented by ectopic expression of *helA* during ICP1^A^ infection, demonstrating that *helA* is necessary for PLE 1 replication.

**Figure 2.**
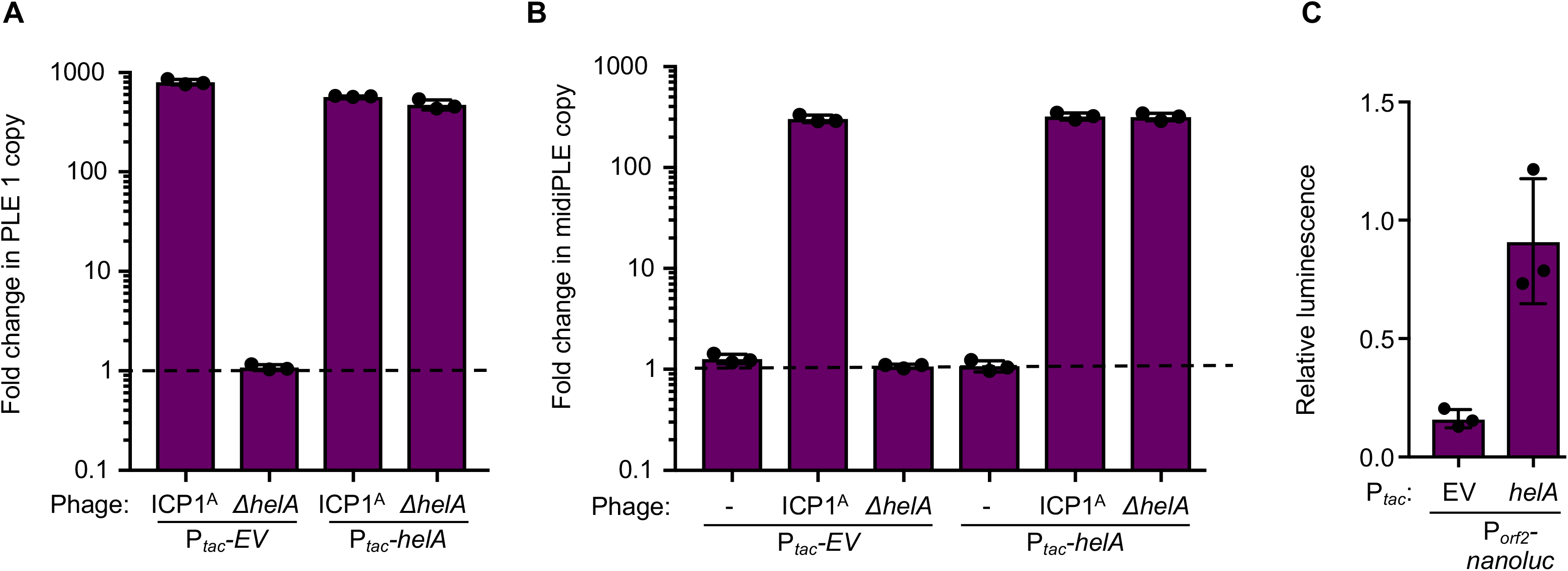
ICP1-encoded *helA* is necessary for PLE replication. **A**, Quantification of change in PLE 1 copy number following infection by the listed ICP1 strain as measured by qPCR. Empty vector (P*_tac_-EV*) and *helA* (P*_tac_-helA*) expression plasmids were induced 20 minutes prior to phage infection. The dashed line indicates no change in copy number. **B**, Quantification of change in midiPLE copy number following infection of midiPLE (+) *V. cholerae ΔlacZ*::P*_tac_-repA* with the listed expression plasmid by the listed ICP1 as measured by qPCR. Ectopic *repA* and expression plasmids were induced 20 minutes prior to phage infection. **C**, Change in luminescence of *P_orf2_-nanoluc* reporter with the listed expression plasmid 20 minutes after infection by ICP1^A^ *ΔhelA* relative to the change in luminescence following infection by ICP1^A^.

SF1B-type helicases are implicated in activities ranging from replication and genome maintenance to transcriptional regulation (Byrd & Raney 2017). Additionally, the S. *aureus* phage parasites, SaPIs, make use of dUTPases as anti-repressors to initiate the transcriptional program of the island, suggesting that these genomic islands can evolve to respond to phage-encoded proteins independent of their biological function for the phage (Tormo-Más et al. 2010; Bowring et al. 2017). As such, we next wanted to determine if *helA* has a direct role in PLE 1 replication or if it is necessary to transcriptionally activate the island to allow for production of PLE 1-encoded proteins, such as *repA*, that are essential for PLE 1 replication. To test the involvement of *helA* in PLE 1 replication, we made use of a minimal PLE replication system referred to as the “midiPLE” (Barth et al. 2019). The midiPLE contains only the endogenous PLE 1 integrase as well as the PLE 1 origin of replication, integrated in the same chromosomal location as PLE 1 in the *V. cholerae* chromosome. This construct is competent to excise from the chromosome following *pexA* expression during ICP1^A^ infection, but is unable to replicate without ectopic expression of the PLE 1-encoded replication initiator, *repA*. When *repA* is provided *in trans*, midiPLE replicates during ICP1^A^ infection (Figure 2B). In comparison to infection with wild-type phage, midiPLE fails to replicate during infection with ICP1^A^ *ΔhelA*. This phenotype can be complemented by expressing *helA in trans*, showing that *helA* is necessary for PLE 1 replication independent of other PLE 1-encoded genes and supporting the conclusion that HelA is directly involved in PLE 1 replication. Interestingly, *helA* is not sufficient to stimulate PLE 1 replication in the absence of ICP1^A^ infection (Figure 2B), indicating that other phage, or possibly *V. cholerae*, components are additionally required to facilitate PLE 1 replication.

### PLE replication contributes to anti-phage gene dosage

In the course of replication sampling during ICP1^A^ infection, we observed a defect in PLE 1-mediated accelerated lysis that correlates with a loss in PLE 1 replication. A culture of PLE (+) *V. cholerae* infected with ICP1 typically lyses 20 minutes after infection, while an infected PLE (−) culture takes upwards of 90 minutes to lyse (O’Hara et al. 2017). However, we observed that cultures infected with ICP1^A^ *ΔhelA* consistently had delays in lysis, suggesting impaired PLE 1 activity, and ectopic expression of *helA* led to intermediate lysis phenotypes (Figure S2A). Though the basis for PLE 1-mediated accelerated lysis is not yet known, we reasoned that robust PLE 1 replication enhances expression of PLE 1-encoded genes merely through increasing the template copy number. To test this hypothesis, we created a nanoluciferase transcriptional reporter cloned downstream of PLE 1 *orf2* (P_*orf2*_*nanoluc*, Figure S2B) to quantify defects in PLE 1 transcription when PLE 1 is unable to replicate. Relative to infection with wild-type ICP1^A^, P_*orf2*_*nanoluc* produced 0.16 times as much luminescence during infection with ICP1^A^ *ΔhelA* (Figure 2C). When PLE 1 replication was restored through ectopic expression of *helA*, the reporter activity resulting from infection with ICP1 *ΔhelA* was restored to wild-type levels, demonstrating that PLE 1 copy number contributes to the global level of PLE 1 transcription. As such, inhibition of PLE 1 replication leads to phenotypes such as delayed lysis during ICP1^A^ infection and potentially contributes to the ability of ICP1^A^ *ΔhelA* to escape PLE 1.

### ICP1 overcomes replication and excision deficient PLE through degradation of the *V. cholerae* chromosome

As ICP1-encoded *pexA* is necessary for PLE 1 excision (McKitterick & Seed 2018) and *helA* is necessary for PLE 1 replication during ICP1 infection (Figure 2A), we next wanted to understand how ICP1^A^ *ΔpexA ΔhelA* is able to overcome PLE 1 (Figure 1B). Even when PLE 1 is challenged by ICP1^A^ *ΔhelA* and is unable to replicate leading to transcriptional deficiencies, PLE 1 is still able to excise from the *V. cholerae* chromosome and is more inhibitory than when it is maintained in the chromosome, leading us to speculate that the position of PLE 1 in the cell, either intra-or extrachromosomal, is important for its activity. Phages are known to encode nucleases that attack the bacterial chromosome, freeing up nucleosides that can then be incorporated into newly synthesized phage genomes (Warner et al. 1970). Additionally, deep sequencing of the total DNA in ICP1 infected *V. cholerae* cells shows that the proportion of reads mapping to the *V. cholerae* chromosomes decreases over the course of infection (Barth et al. 2019). This observation led us to hypothesize that nucleolytic activity encoded by ICP1^A^, deployed to degrade the *V. cholerae* chromosome during infection, is able to degrade PLE 1 when PLE 1 is stuck in the chromosome unable to replicate, allowing for ICP1^A^ to form some small plaques on PLE (+) *V. cholerae*. To test this hypothesis, we made use of a minimal PLE excision system, the miniPLE, that has the PLE 1-encoded integrase but lacks an origin of replication (Figure 3A). Thus during infection, the miniPLE excises from the host chromosome and circularizes, but does not replicate (McKitterick & Seed 2018). To simulate an excision-deficient miniPLE, we created miniPLE_CD_, which possesses a point mutation in the catalytic serine residue in the miniPLE-encoded integrase, making the integrase catalytically dead and rendering the construct unable to excise from the chromosome (Figure 3B).

**Figure 3.**
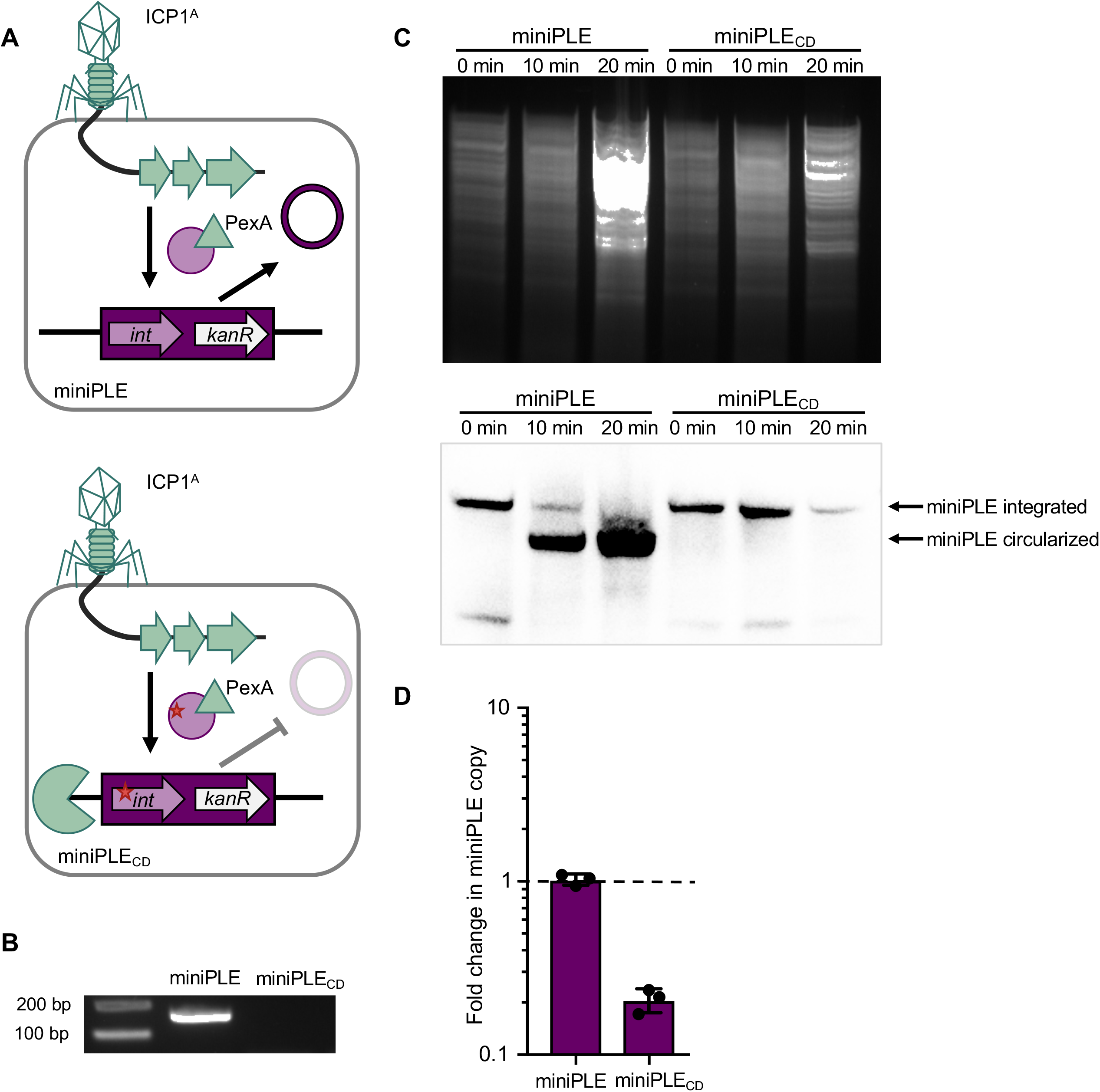
Excision and replication deficient PLE is susceptible to ICP1-mediated chromosomal degradation. **A**, Cartoon of miniPLE during ICP1 infection. Top, miniPLE-encoded Int (circle) is directed to excise miniPLE during ICP1 infection by ICP1-encoded PexA (triangle), leading to a single-copy circularized miniPLE episome. Bottom, catalytically dead miniPLE_CD_ Int (circle with red star) is unable to excise miniPLE during ICP1 infection, potentially rendering the miniPLE susceptible to phage-mediated chromosomal degradation (pac-man). **B**, Circularization PCR of the miniPLE indicated from boiled ICP1^A^ plaques on the host indicated. **C**, (Top) Total DNA prepped from equal numbers of miniPLE or miniPLE_CD_ cells infected by ICP1^A^ at the listed timepoints and imaged via Southern blot (bottom) with a probe against the miniPLE *kanR* cassette. **D**, Change in copy number of the miniPLE indicated 30 minutes following ICP1^A^ infection as measured by qPCR.

Total DNA from ICP1^A^ miniPLE and miniPLE_CD_ infected cells was digested, run on an agarose gel, and the stability of the miniPLE was observed via Southern blot (Figure 3C). During the course of ICP1^A^ infection, the miniPLE successfully excises from the *V. cholerae* chromosome and is maintained as an episome. Conversely, the amount of miniPLE_CD_, which is unable to excise from the chromosome, decreases by 20 minutes following ICP1^A^ infection (Figure 3C, bottom), relative to the amount of total DNA prepped from the cells (Figure 3C, top), suggesting that the copy number of miniPLE_CD_ decreases as a result of ICP1^A^ infection. Quantification of miniPLE via qPCR further demonstrates that the excision-competent miniPLE is maintained as a stable episome with no change in copy number during ICP1^A^ infection (Figure 3D). In comparison, the miniPLE_CD_ that is unable to escape the *V. cholerae* host chromosome decreased in copy number during infection with ICP1^A^, indicating that it is susceptible to ICP1^A^-mediated chromosomal degradation. Thus, not only is PLE mobilization important for HGT (O’Hara et al. 2017; Barth et al. 2019), but it is also essential for PLE escape from ICP1 takeover of the *V. cholerae* host.

### Diverse SF1B helicases are maintained in ICP1 and contribute to ICP1 fitness

Due to the importance of PLE replication in PLE gene dosage and avoiding ICP1-mediated host takeover, we next hypothesized that ICP1 would evolve to abolish PLE replication by accumulating mutations in the *helA* allele, indicative of co-evolution between the two entities. To identify signatures of co-evolution, we examined HelA from sequenced isolates of ICP1 that had been recovered from epidemic sampling in Dhaka, Bangladesh. HelA from ICP1 isolated from epidemic sampling from 2001 to 2017 is over 99% identical at the amino acid level indicating that there is either little pressure for HelA to evolve over time, or that HelA mutations cannot be tolerated in nature (Table S6). Though there is no change in the ability of ICP1 to replicate in a single round of infection in the absence of *helA* (Figure 1E), ICP1^A^ *ΔhelA* forms plaques that are on average 0.75 times smaller than wild-type phage plaques (Figure 4A). This size defect indicates that mutant phage are less fit in the absence of *helA* and supports the notion that functional *helA* must be maintained by ICP1 in nature.

**Figure 4.**
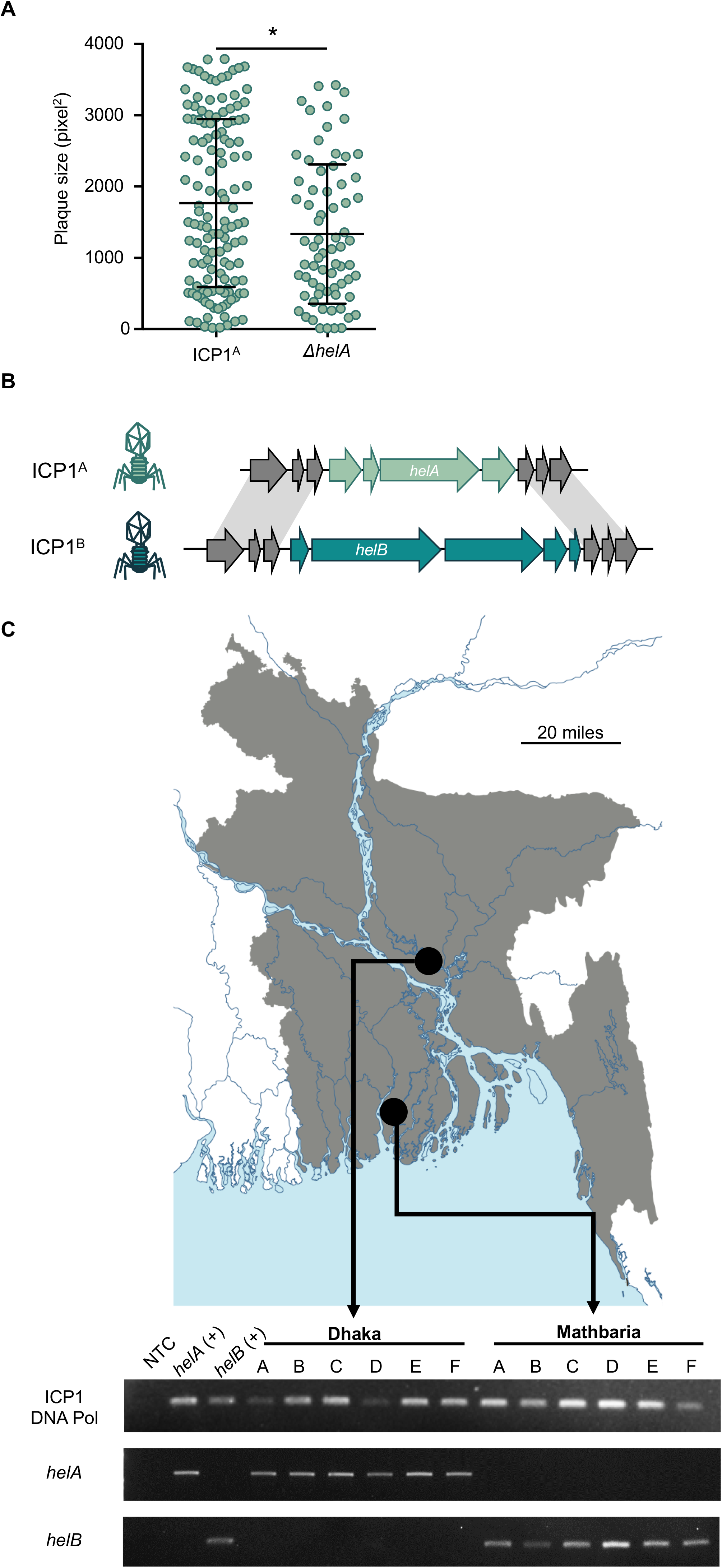
ICP1 encodes one of two accessory helicase alleles. **A**, Plaque size of listed phage on PLE (−) *V. cholerae*. *p<0.01. **B**, Cartoon of ICP1 accessory helicase locus. Grey arrows indicate gene products shared between the two phages, while the mint arrows indicate gene products unique to the *helA* locus and turquoise arrows indicate gene products unique to the *helB* locus. **C**, Map (Vecteezy 2019) of distribution of SF1B-type helicases alleles in ICP1 isolates shed by cholera patients in Bangladesh. Top, map of Bangladesh with Dhaka and Mathbaria marked. Bottom, agarose gel showing PCR detection of the conserved DNA polymerase (*gp58*), *helA*, and *helB* in ICP1 isolates from cholera patient stools collected in Dhaka or Mathbaria. Phage isolates are listed in Table S8.

Despite having a high degree of conservation, *helA* is not considered part of the core ICP1 genome (Angermeyer et al. 2018): two phage isolates recovered from cholera patient stool samples from Dhaka in 2006 do not encode *helA*, but instead have an alternative SF1B-type helicase in the same locus, which we call *<u>heli</u>case <u>B</u>* (*helB*) (Figure 4B). HelB is 24% identical to HelA, with a conserved P-loop ATPase domain, but HelB has an extended C-terminus that contains a domain of unknown function, DUF2493 (Figure S3A). In addition to having low sequence identity, *helA* and *helB* are flanked by different, unrelated genes each encoding products with no predicted structure or function (Figure 4B), suggesting that while ICP1 is unable to lose *helA* in nature in an attempt to avoid hijacking by PLE for replication, ICP1 may swap *helA* for a distinct accessory helicase.

We then performed a BLASTP search of the National Center for Biotechnology Information’s nonredundant protein sequence database to identify the origin of *helA* and *helB*. Homologs of HelA are commonly found in phages of marine bacteria, and, particularly, in a group of related myoviruses that infect non-cholera *Vibrios* (Figure S3B). Of note, two of the *Vibrio* phages were also predicted to encode a homolog of one of the proteins flanking HelA in ICP1^A^, indicating that the *helA* locus could have been shared with a common ancestor of these phages. Conversely, HelB is more divergent, with the only identifiable homolog found in a *Pseudoalteromonas* phage that is also predicted to have the same DUF2493 C-terminus. These HelB proteins cluster on a more distant branch than the HelA homologs (Figure S3B), supporting the hypothesis that *helB* was horizontally acquired by ICP1. Altogether, SF1B helicases are readily found in marine phages, and ICP1 encoding *helA* are the dominant ICP1 shed by cholera patients in Dhaka between 2001-2017.

Most epidemic sampling of ICP1 from cholera patients has been done in the urban cholera endemic site in Dhaka; however, we recently began sampling cholera patients at a rural and estuarine site in Mathbaria, Bangladesh. In contrast to what was observed in ICP1 isolates from Dhaka in the 2017 epidemic period, all the ICP1 isolates recovered from cholera patients in Mathbaria encoded the *helB* allele (Figure 4C). One representative isolate from Mathbaria from 2017, referred to here as ICP1^B^, is over 99.8% identical to ICP1^A^ across 90% of the genome, with 205 of 227 ICP1^B^ predicted open reading frames being shared with ICP1^A^. The resurgence and dominance of *helB* in the Mathbaria epidemic sampling suggests that there could be a selective advantage for ICP1 encoding *helB* rather than *helA* in this region.

As ICP1^B^ is not isogenic to ICP1^A^, we first wanted to characterize the role of *helB* in ICP1^B^ fitness. Similar to HelA, HelB is detectable by Western blot within 8 minutes of infection (Figure 5A), again coinciding with ICP1 replication (Barth et al. 2019). Also similar to *helA, helB* is not essential for ICP1^B^, and ICP1^B^ *ΔhelB* is able to form plaques in the absence and presence of PLE 1 (Figure 5B). Interestingly, ICP1^B^ *ΔhelB* forms plaques on PLE (+) *V. cholerae* with a higher efficiency than ICP1^A^ *ΔhelA*, suggesting that ICP1^B^ has evolved other ways to limit PLE-mediated anti-phage activity.

**Figure 5.**
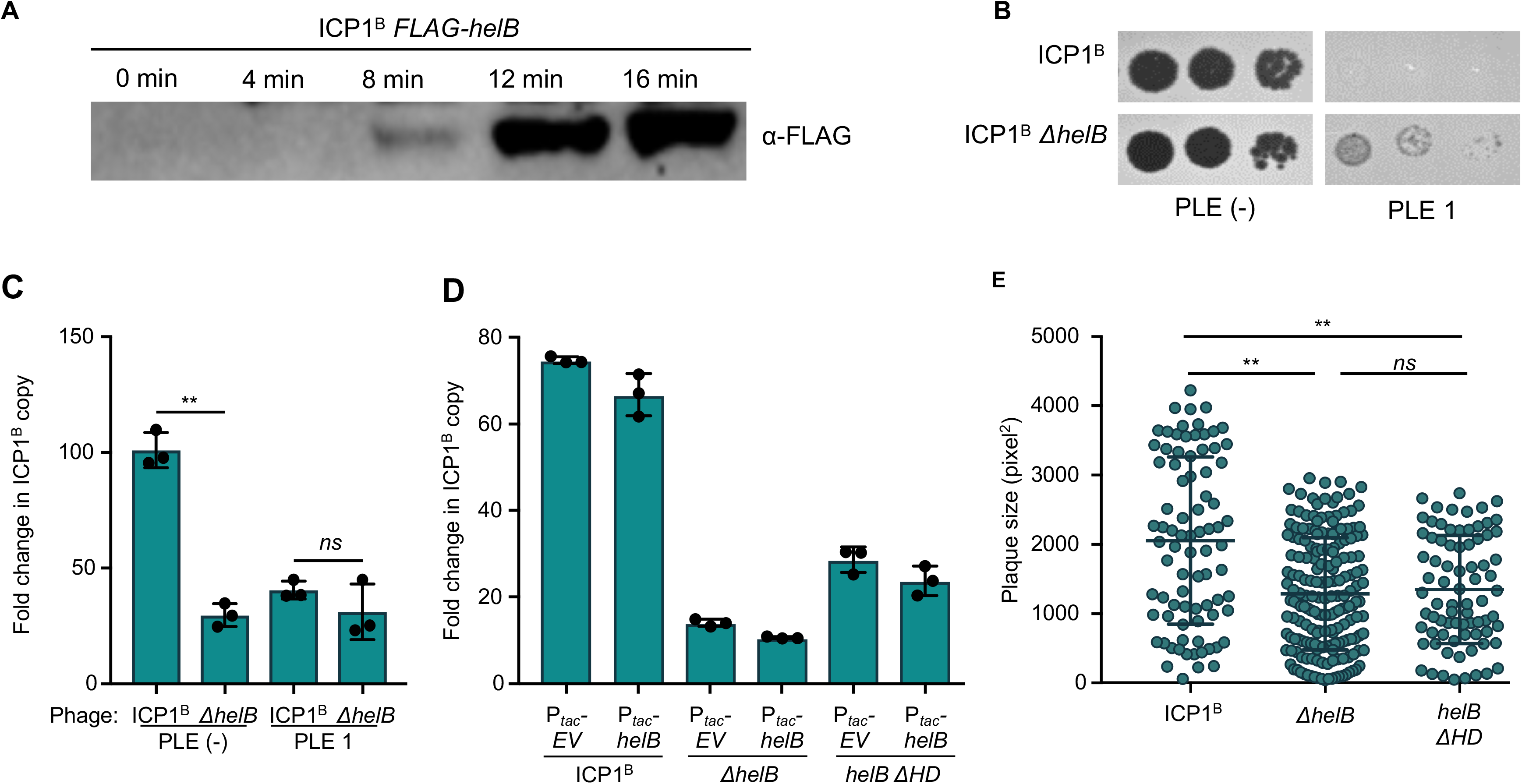
Loss of *helB* permits escape from PLE but leads to a defect in ICP1 fitness. **A**, Western blot of endogenously FLAG-tagged *helB* at the listed time points following infection of PLE (−) *V. cholerae*. **B**, Tenfold dilutions of ICP1 spotted on the listed *V. cholerae* lawns. **C**, Fold change in ICP1 copy number following 20 minutes of infection of the listed *V. cholerae* host as measured by qPCR. **D**, Fold change in ICP1 copy number following 20 minutes of infection of the listed *V. cholerae* host as measured by qPCR. Ectopic expression was induced 20 minutes prior to phage infection. **E**, Plaque size of listed phage on PLE (−) *V. cholerae*. **p<0.001, *ns* not significant.

We next wanted to see if ICP1^B^ replication was impacted by the *helB* deletion. In contrast to *ΔhelA* in ICP1^A^, ICP1^B^ *ΔhelB* is significantly impaired for replication during the course of infection compared to wild-type ICP1^B^ (Figure 5C), indicating that although *helB* is not necessary for ICP1^B^ replication, it does have a more central role in phage fitness. Consistent with the observation that PLE 1 decreases the ability of ICP1^A^ to replicate (Figure 1E), replication of ICP1^B^, too, is impacted negatively by PLE 1; however, ICP1^B^ *ΔhelB* does not restore the ability of ICP1^B^ to replicate in the presence of PLE 1 (Figure 5C), demonstrating a more severe fitness effect associated with losing the accessory helicase on ICP1^B^ than on ICP1^A^ independent of the presence of PLE 1.

To confirm the role of *helB* in diminished ICP1^B^ fitness, we next ectopically expressed *helB* to complement the mutant phage. However, we could not complement the replication defect for ICP1^B^ *ΔhelB* by ectopic expression of *helB*, suggesting that the observed decrease in ICP1 fitness may not be due to direct loss of the *helB* gene product (Figure 5D). To minimize potential polar effects of *ΔhelB*, a targeted mutation was made to remove 25 amino acids encompassing the helicase domain (HD) that contains the Walker A motif necessary for ATP hydrolysis (Blair et al. 2009). While ICP1^B^ *helB ΔHD* had increased phage replication relative to the clean *helB* deletion, there was still a defect in replication that could not be complemented (Figure 5D), suggesting that ectoptic expression may not be able to achieve the appropriate timing or dosage of *helB* expression, or that the fitness cost is not a result of loss of HelB *per se*. Due to the complex nature of phage genomes and tight regulation of phage gene expression, disruption of even the HD domain of *helB* could have detrimental effects on uncharacterized *in cis* sites that could contribute to poor fitness. The fitness defect associated with mutant *helB* was also observed as a decrease in plaque size, with both ICP1^B^ *ΔhelB* and ICP1^B^ *helB ΔHD* forming plaques that are, on average, less than 0.66 times the size of ICP1^B^ (Figure 5E). Altogether, ICP1^B^ is less fit in the absence of *helB*, consistent with the observation that all natural ICP1 isolates encode one of two SF1B-type helicases, either *helA* or *helB*.

### PLE exploits phage-encoded distinct SF1B-type helicases to drive replication during ICP1 infection

Given that PLE 1 replication requires *helA* (Figure 2A), and ICP1 with *helB* are dominant in Mathbaria, we were tempted by the possibility that phage with *helB* could be selected for as a mechanism to impede PLE 1 replication during infection. Hence, we next assessed if *helB* could also support PLE 1 replication. Consistent with the inverse relationship between ICP1 and PLE 1 replication, PLE 1 still replicated when infected with ICP1^B^, and as with *ΔhelA*, PLE 1 replication was not observed in the absence *helB* (Figure 6A), indicating that *helB* is also necessary for PLE 1 replication despite HelB having less than shared 25% shared amino acid identity with HelA (Figure S3A). Further, ectopic expression of *helB* complemented the defect in PLE 1 replication observed during infection with ICP1^B^ *ΔhelB*, and ectopic expression of *helA* was likewise sufficient to restore PLE 1 replication during infection with ICP1^B^ *ΔhelB* (Figure 6A). These data demonstrate that PLE 1 is able to harness either ICP1-encoded accessory helicase independent of the ICP1 isolate that is infecting the host. Additionally, the shared ability of these non-isogenic ICP1 isolates to drive PLE 1 replication implicates functionally conserved gene products in ICP1 isolates, in addition to *helA* and *helB*, that are required for PLE 1 replication.

**Figure 6.**
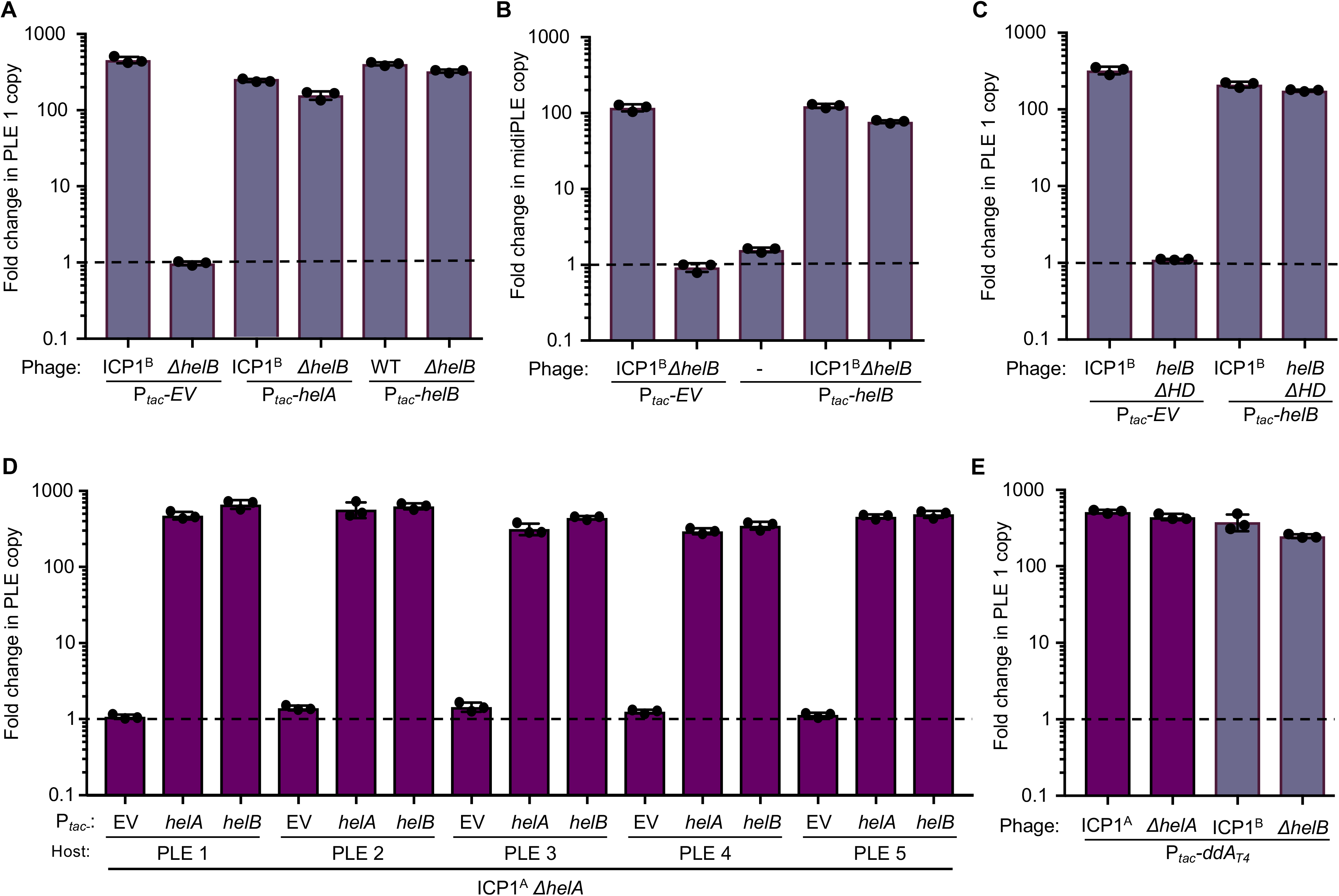
PLEs can exploit unrelated phage-encoded SF1B-type helicases for replication. Replication of PLE 1 (**A,C**) or midiPLE (**B**) 20 minutes following infection of *V. cholerae* with the listed expression vectors by the listed ICP1^B^ variant as measured by qPCR. Vectors were induced 20 minutes prior to infection. Dashed line indicates no change in copy. **D**, Replication of the listed PLE in an isogenic *V. cholerae* background 20 minutes following infection by ICP1^A^ *ΔhelA*. Ectopic vectors were induced 20 minutes prior to infection. Dashed line indicates no change in copy. **E**, Replication of PLE 1 20 minutes following infection by the listed phage as measured by qPCR. Ectopic expression of *dda* from *E. coli* phage T4 was induced 20 minutes prior to infection. Dashed line indicates no change in copy.

Similar to *helA*, we next used ICP1^B^ *ΔhelB* to probe for midiPLE replication following ectopic expression of *repA*. As expected, midiPLE replicated when infected with ICP1^B^ but failed to replicate in the absence of *helB*, indicating that *helB* is also directly involved in PLE 1 replication (Figure 6B). Like *helA, helB* is also not sufficient to stimulate PLE 1 replication in the absence of ICP1^B^, showing that PLE 1 is still dependent on additional replication machinery from ICP1^B^. We additionally confirmed that the ability of HelB to hydrolyze ATP is required for HelB to facilitate PLE 1 replication by testing the ICP1^B^ *helB ΔHD* variant, and, as anticipated, the helicase activity of *helB* is necessary for PLE 1 replication (Figure 6C).

The first ICP1 isolate identified with the *helB* allele was from Dhaka in 2006 when PLE 2 *V. cholerae* were being shed by cholera patients (O’Hara et al. 2017; McKitterick et al. 2019), leading us to evaluate if the two helicase alleles have different capacities to facilitate replication of different PLEs during infection with ICP1. To test this hypothesis, we first infected isogenic *V. cholerae* harboring each of the five characterized PLEs with ICP1^A^ and observed that all PLEs replicated equally well (Figure S4). Next, we determined that *helA* is necessary for replication of all five PLEs during ICP1^A^ infection and that replication can be complemented with ectopic expression of *helA* (Figure 6D). To evaluate if each PLE can additionally use *helB* to support replication, we also complemented ICP1^A^*ΔhelA* with ectopic expression of *helB* and found that in fact all five PLEs can use either one of the two ICP1-encoded accessory helicases for replication.

As current data supports the model that PLE responds specifically to ICP1 infection (O’Hara et al. 2017; McKitterick & Seed 2018), we next wanted to determine if PLE’s capacity to exploit either *helA* or *helB* to drive PLE replication is specific to ICP1-encoded proteins or in general to SF1B-type helicases. To address the specificity of the interaction, we ectopically expressed the SF1B-type helicase *dda from E. coli* phage T4 during infection with either ICP1^A^ *ΔhelA* or ICP1^B^ *ΔhelB*. T4 Dda is only 16% identical to either HelA or HelB and does not group with the marine phage SF1B-type helicases (Figure S3B). Although PLE 1 cannot replicate while infected with either of these *Δhel* phage alone (Figures 6A and 6D), expression of *dda* was sufficient to support PLE 1 replication in the absence of ICP1-encoded accessory helicases (Figure 6E). Despite the apparent specificity between PLE and ICP1, the ability of PLE to exploit a variety of phage-encoded accessory helicases reveals flexibility in at least one requirement for PLE replication, and suggests that swapping of helicase alleles by ICP1 isolates is not a beneficial strategy to mitigate PLE parasitism.

## Discussion

In order to defend against viral infection, host resistance mechanisms must have ways by which they prevent or bypass virus mediated host takeover. Eukaryotic DNA and RNA viruses broadly use virally encoded ribonucleases to globally degrade host transcripts in the infected cell, which sabotage their hosts through modulation of transcript and protein levels. This decrease in transcript abundance leads to a downregulation of innate immune responses, processes which are detrimental to the host but are ultimately reversable (Crow et al. 2016; Rivas et al. 2016). Conversely, degradation of the host chromosome is a host takeover process that is unique to phages. Host chromosome degradation has a twofold benefit for the predatory phages: it cleaves and releases nucleosides that can be incorporated into the rapidly replicating phage genome, and it can also destroy the template needed for expression of anti-phage genes encoded by the bacterial host. With the imminent shutdown of the host upon phage infection, it is not surprising to find that many bacterial defense systems, such as restriction-modification and some CRISPR-Cas systems, are expressed constitutively. By balancing the ability to discern between self and non-self, these systems are safely deployed in the absence of infection. Conversely, more self-destructive defense mechanisms, such as toxin/antitoxin systems and abortive infection systems cannot be constitutively active due to lethal outcomes and must be induced upon infection. Thus, for an inducible defense system like PLE, and perhaps many phage parasites, mobilization to evade host shutdown is critical. When PLE is unable to replicate or excise from the chromosome, it no longer fully blocks plaque formation by ICP1 and is susceptible to ICP1-mediated degradation of *V. cholerae’s* chromosomes.

As a defense island and phage parasite of ICP1, the *V. cholerae* PLE has become highly evolved to make use of phage-encoded gene products to drive its anti-phage program (McKitterick & Seed 2018). Here, we characterize a new ICP1-PLE interaction: PLE hijacks a non-essential ICP1-encoded SF1B-type helicase to drive PLE replication during infection, making PLE the first characterized phage satellite that makes use of replication machinery encoded by its helper phage. In comparison, the well-studied SaPIs make use of their bacterial host’s replication machinery and are able to autonomously replicate in the absence of helper phage (Úbeda et al. 2008). PLE’s unique requirement for the phage-encoded helicase also underscores the differences between the helper phages that induce these chromosomal islands, with PLE being induced by a lytic phage that encodes its own replication machinery and SaPIs being induced by an activated temperate phage that also exploits its host-encoded replication machinery (Úbeda et al. 2008). The fact that *helA* expression alone is not sufficient to drive midiPLE replication in the absence of ICP1 infection implicates other ICP1-encoded replication proteins in facilitating PLE replication. Aside from the SF1B-type helicases, the potential role for ICP1’s replication machinery in PLE replication remains to be elucidated. As T4 *dda* has been observed to have a role in T4 origin initiation during origin-dependent replication (Brister 2008), we speculate that *helA* has a similar role in facilitating origin firing in PLE by interacting with a conserved part of the PLE machinery and recruiting a conserved ICP1 replication protein. ICP1 is predicted to encode a DNA polymerase and primase/helicase reminiscent of machinery that drives the *E. coli* phage T7 replisome (Barth et al. 2019). Further work remains to identify what roles, if any, these replisome proteins have in PLE mobilization.

Despite not being essential, all ICP1 isolates encode an accessory SF1B-type helicase, as do several marine phages (Figure S3B) (Kauffman et al. 2018). Of note, one of these marine *Vibrio* phages is also predicted to encode a complete Type 1-F CRISPR-Cas system, which is of the same type that is encoded by some isolates of ICP1 to target and overcome PLE activity (Seed et al. 2013; McKitterick et al. 2019), suggesting that ICP1 could be exchanging genetic material with or could be related to these marine phages infecting non-cholera *Vibrios*. The fitness costs associated with losing the accessory SF1B-type helicase, as measured by plaque size, implicate both *helA* and *helB* in maintaining optimal phage fitness, though the precise role for these accessory helicases in the phage lifecycle remains to be determined. The ease with which PLE is able to make use of ectopically expressed *helB* compared to the inability of ectopically expressed *helB* to complement the ICP1^B^ *ΔhelB* replication deficiency suggests that these helicases play a specialized role in the phage lifecycle that is more complex than for PLE.

Due to the variability between ICP1 isolates, we see that PLE has evolved to make use of not just two unrelated ICP1-encoded helicases, but also of T4 *dda*, the prototypical but unrelated SF1B helicase. The ability of PLE to replicate using several dissimilar helicases implicates strong evolutionary pressures for maintenance of PLE replication in response to ICP1 evolution. While *helA* is only one of the seemingly wide variety of ICP1 inputs that contribute to PLE activity, SaPIs have similarly evolved to overcome variability in helper phage induction cues (Bowring et al. 2017). The apparent promiscuity of the SaPI master repressor allows for recognition of structurally dissimilar but functionally conserved phage proteins to ensure SaPI excision, replication and spread, despite their helper phage’s attempts to avoid SaPI induction.

It is also imperative for PLE to be able to make use of either one of the ICP1-encoded helicases to continue PLE propagation through epidemic *V. cholerae* populations, thus selecting for PLE genes that are able to make use of dissimilar helicases despite the capacity of ICP1 to swap one helicase for another.

The striking spatial separation between the ICP1^A^ and ICP1^B^ populations that were shed by cholera patients in Bangladesh during the same epidemic period suggests that slight variations in the phage strain, such as the difference between *helA* and *helB*, can have large differences in the makeup of phage populations. Indeed, the ability of ICP1^B^ *ΔhelB* to form plaques in the presence of PLE (Figure 5B) suggests that ICP1^B^ should dominate in the presence of PLE (+) *V. cholerae;* however, the greater fitness cost for ICP1^B^ in the absence of *helB*, as evidenced by the diminished ability of ICP1^B^ *ΔhelB* to replicate, suggests that the loss of *helB* in the presence of PLE (+) *V. cholerae* is ultimately detrimental to the phage population.

The necessity of excision and replication of PLE during ICP1 infection highlights a crucial role for mobilization of inducible phage defense systems during phage infection. In order for inducible defenses to functionally protect a host cell from phage infection, they must be able to overcome the infecting phage’s destruction of the host chromosome. Elements independent of the host chromosome, such as plasmids, seem to be somewhat protected from degradation by lytic phages (Keen et al. 2017). It thus stands to reason that the observed high prevalence of phage defense systems encoded on genomic islands (Makarova et al. 2011) may be in part due to the ability of genomic islands to mobilize during infection and escape phage-mediated host takeover, with the potential of horizontal transfer or the ability to escape from a dying host as an added benefit. Through experimental and *in silico* validation, more phage defense islands have been identified and characterized, albeit often in a context independent from infection by a native phage. Given the propensity of some phages to degrade their host chromosome during infection and the need for protective MGEs to escape host takeover, it will be interesting to further explore if other inducible defense islands mobilize in response to phage infection and are in fact unrecognized phage satellites.

## ACKNOWLEDGMENTS

This research was funded by the National Institute of Allergy and Infectious Diseases grant number R01AI127652 (K.D.S.). A.C.M. received support from the Kathleen L. Miller Fellowship from the Henry Wheeler Center for Emerging and Neglected Diseases. K.D.S. is a Chan Zuckerberg Biohub Investigator. The authors are especially thankful to icddr,b hospital and laboratory staff for their support, in particular Shirajum Monira, Fatema-tuz Johura, Marzia Sultana, Kazi Zillur Rahman, and Monika Sultana. M.A. of icddr,b, thanks the governments of Bangladesh, Canada, Sweden, and United Kingdom for providing core/unrestricted support. We also wish to thank members of the Seed Lab for helpful discussions, and Angus Angermeyer, specifically, for help with the annotation/computational analyses.

## AUTHOR CONTRIBUTIONS

Conceptualization, A.C.M and K.D.S.; Investigation, A.C.M. and S.G.H.; Resources, M.A.; Writing – Original Draft, A.C.M. and K.D.S.; Writing – Review & Editing, A.C.M, S.G.H, M.A. and K.D.S.; Funding Acquisition, A.C.M., M.A. and K.D.S.

## Supplementary Figures

**Figure S1. Related to Figure 1. ICP1^A^ *ΔpexA ΔhelA* does not accumulate mutations to escape PLE**. Three sets of plaques (a-c) from ICP1 *ΔpexA ΔhelA* on PLE 1 *V. cholerae* were picked and the efficiency of plaquing on PLE 1 relative to PLE (−) *V. cholerae* was tested.

**Figure S2. Related to Figure 2. Infected PLE *1 V. cholerae* demonstrate altered lysis kinetics when PLE 1 does not replicate**. **A**, OD_600_ of the PLE 1 *V. cholerae* over time after infection with the listed ICP1. **B**, Cartoon of the PLE 1-encoded nanoluciferase reporter with nanoluciferase (nL) encoded downstream of PLE 1 *orf2*. Line break in the PLE genome is shown for simplicity and no other mutations are present.

**Figure S3. Related to Figure 4. SF1B-type helicases are found in a variety of marine phages**. **A**, Praline alignment (Bawono & Heringa 2014) of ICP1^A^ HelA, ICP1^B^ HelB, and T4 Dda. The Walker A motif used in ATP hydrolysis is indicated by asterisks, and the Duf2493 in HelB is underlined. **B**, Phylogenetic analysis of SF1B-type helicases, phages used are listed in Table S7.

**Figure S4. Related to Figure 6. PLE replication is not altered by ectopic expression of *helA* or *helB***. Replication of the listed PLE in an isogenic *V. cholerae* background 20 minutes following infection by ICP1^A^. Ectopic vectors were induced 20 minutes prior to infection. Dashed line indicates no change in copy.

## STAR Methods

### CONTACT FOR REAGENT AND RESOURCE SHARING

Further information and requests for resources and reagents should be directed to and will be fulfilled by the lead Contact Kimberley Seed.

### EXPERIMENTAL MODEL AND SUBJECT DETAILS

#### Bacterial Growth Conditions

The bacterial strains and plasmids used in this study are listed in Tables S2 and S5. All bacterial strains were grown at 37°C in LB with aeration or on LB agar plates. The following antibiotics were used as necessary: streptomycin (100 μg/mL), spectinomycin (100 μg/mL), kanamycin (75 μg/mL), ampicillin, (V. *cholerae* 50 μg/mL, *E. coli* 100 μg/mL), chloramphenicol (V. *cholerae* 1.25 μg/mL, *E. coli* 25 μg/mL). Ectopic expression constructs in *V. cholerae* were induced 20 minutes prior to ICP1 infection with 1 mM Isopropyl β-D-1-thiogalactopyranoside (IPTG) and 1.5 mM theophylline.

#### Phage Growth Conditions

The phage isolates used in this study are listed in Table S3. Phage were propagated using the soft agar overly method and high titer stocks were made by polyethylene glycerol precipitation and stored in sodium chloride-tris-EDTA (STE) buffer (Clokie & Kropinski 2009).

#### Phage isolation from rice water stool

The collection of cholera patient rice water stool (RWS) was approved by the icddr,b institutional review board. All samples were deidentified and written informed consent was obtained from adult participants and from the guardians of children. Stool samples were mixed with glycerol in cryovials, frozen, until being processed at the University of California, Berkeley. For processing, samples were thawed and grown on thiosulfate-citrate-bile salts-sucrose agar, or used to inoculate alkaline peptone water (APW) for outgrowth. Liquid APW cultures were struck out on agar plates and aliquots were frozen with glycerol. Individual colonies selected from plates were confirmed as *V. cholerae* by PCR. These isolates of *V. cholerae*, in addition to the PLE (−) laboratory strain, were used to isolate phages directly from the RWS glycerol stocks and from frozen APW outgrowths. Isolated phages were plaque purified twice after isolation.

### METHOD DETAILS

#### Bacterial and phage cloning conditions

Bacterial mutants were cloned using SOE (splicing by overlap extension) PCR and introduced by natural transformation (Dalia et al. 2014). Plasmids were constructed using Gibson Assembly or Golden Gate Assembly. Phage mutants were constructed using CRISPR-Cas engineering as previously described (Box et al. 2016; McKitterick & Seed 2018). Briefly, an editing template with the desired deletion was cloned into a plasmid and *V. cholerae* harboring this plasmid was infected by the ICP1 strain of interest. Ten plaques of the passaged phage were collected and mutants were selected on *V. cholerae* engineered to encode an inducible Type 1-E CRISPR-Cas system and a plasmid with a spacer targeting the gene of interest. Mutant phages were verified via Sanger sequencing and purified two times on the targeting host before storing in STE. Total phage gDNA was prepped with a DNeasy Blood & Tissue Kit (Qiagen).

#### Phage plaquing conditions

Spot plates were performed as before (McKitterick & Seed 2018). Briefly, mid-log *V. cholerae* was added to 0.5% molten LB agar poured on a solid agar plate and allowed to solidify. Ten-fold dilutions of phage were applied to the surface in 3 μL spots and allowed to dry. Plates were incubated at 37°C. Images are representative of at least two independent experiments. The efficiency of plaquing (EOP) was calculated by comparing the number of plaques a given phage forms on PLE (−) *V. cholerae* relative to the number of plaques formed on PLE (+) *V. cholerae*. Each EOP was calculated in triplicate, and the limit of detection is the point at which the phage is unable to productively infect the PLE (+) host while still forming plaques on a PLE (−) host. Plaque size was determined by imaging and quantifying with ImageJ at least 20 plaques each from 3 independent replicates in 0.5% agar overlay on PLE (−) *V. cholerae*. Significance was determined through a nonparametric T Test.

#### qPCR conditions

Fold change in genome copy was performed as before (O’Hara et al. 2017) with slight modification. Fold change in ICP1 copy number was measured by growing cells to an OD_600_ of 0.3, infected with a multiplicity of infection (MOI) of 0.01, and a sample was boiled for 10 minutes as a starting value. Infected cells were returned to the incubator for 20 minutes, at which point another sample was taken and boiled. Boiled samples were diluted 1:50 and used as a template in the qPCR reaction. To measure the fold change in PLE and midiPLE copy number, cells were grown to an OD_600_ = 0.3 and the initial sample was immediately taken prior to addition of phage at an MOI of 2.5 and boiled for 10 minutes. Samples were taken at 20 minutes after infection, boiled for 10 minutes, and diluted 1:1000. Quantification of the fold change in miniPLE copy was measured during infection with an MOI of 5, with samples taken immediately prior to infection and 30 minutes after infection and boiled for 10 minutes. Boiled samples were diluted 1:100 and used as template. Experiments with ectopic expression constructs were induced at OD_600_ = 0.2 for 20 minutes and then normalized to OD_600_ = 0.3 prior to infection. All samples were run in biological triplicates and technical duplicates. The template was mixed with the primers listed in Table S4 and IQ Sybr Green Master Mix (Bio-rad) and run on a CFX Connect Real-Time PCR Detection System (Bio-rad). Fold change was measured as the amount of DNA in the sample at 20 or 30 minutes after infection relative to the amount of DNA in the sample at T=0. Significance was measured by 2-tailed T Test.

#### Western Blots

PLE (−) *V. cholerae* was grown to an OD_600_ = 0.3 and infected with the endogenously FLAG-tagged ICP1 listed. At the listed timepoints, 1 mL samples were collected and mixed with equal volume ice-cold methanol and centrifuged at 13000 rpm for 3 minutes at 4°C. Pellets were washed with ice-cold PBS, resuspended in 1x Laemmli buffer, and boiled for 10 minutes at 99°C. Total protein was run on a 10% Stain Free TGX SDS-PAGE gel (Bio-rad). Primary Rabbit-α-FLAG antibodies (Sigma) were used at a dilution of 1:5000 and detected with goat-α-rabbit-HRP conjugated secondary antibodies at a dilution of 1:5000 (Bio-rad). Clarity Western ECL Substrate (Bio-rad) was used to develop the blots and a Chemidoc XRS Imaging System (Bio-rad) was used to image.

#### Lysis kinetics and Nanoluciferase assay

PLE (+) *V. cholerae* cells were grown to an OD_600_ = 0.2 and the listed ectopic expression constructs were induced for 20 minutes. Cells were then normalized to an OD_600_ = 0.3 and infected at an MOI of 2.5. For lysis kinetics, OD_600_ was monitored for 30 minutes. For nanoluciferase, 100 μL cells were sampled at T=0 and T=20 minutes after infection and added to 100 μL ice cold methanol. Luminescence was measured in a Spectra Max i3x plate reader (Molecular Devices) using the Nano-Glo Luciferase Assay System (Promega). Relative luminescence was calculated by dividing the luminescence detected after infection with the knockout phage relative to the luminescence detected after infection with the WT phage.

#### PCR conditions

PLE circularization PCRs were performed as described (McKitterick & Seed 2018). Briefly, plaques on the miniPLE or miniPLE_CD_ hosts were picked into 50 μL of water and boiled for 10 minutes. Boiled template (2 μL) was used with the primers listed in Table S4 to detect miniPLE circularization. Detection of ICP1 *DNA pol, helA*, and *helB* from ICP1 isolates were performed on 5 – 30 ng prepped gDNA from isolated phage with the primers listed in Table S4. PCRs were run on 2% agarose gels and visualized with GelGreen.

#### Southern Blots

A probe against miniPLE was created using the DIG-High Prime DNA Labeling and Detection Started Kit I (Sigma). Cells were grown up to OD_600_ = 0.3 with kanamycin and infected with ICP1^A^ at an MOI of 5. At the timepoints indicated, 5 mL of cells were harvested and mixed with 5 mL ice cold methanol. Samples were spun at 7000xg at 4°C for 5 minutes. Pellets were washed ice cold PBS and spun again. Total DNA was extracted from the pellets with the DNeasy Blood & Tissue Kit (Qiagen). Equal volumes of samples (between 1.5 and 4.1 μg DNA) were digested overnight with EcoRV-HF and SalI-HF (NEB) and run on a 0.7% agarose gel and visualized with GelRed. The agarose gel was washed briefly and incubated with 0.25 N HCl for 15 minutes, washed again, denatured in 0.4 M NaOH for 20 minutes, and transferred overnight. DNA was fixed by baking the blot at 120 °C for 30 minutes, and hybridized with 17 ng/mL miniPLE probe overnight at 42°C. The blot was detected with the DIG-High Prime DNA Labeling and Detection Started Kit I (Sigma) and CSPD™ Substrate (ThermoScientific) and visualized on a Chemidoc XRS Imaging System (Bio-rad).

#### Computational analyses

Escape ICP1 *ΔpexA* phage were isolated from PLE (+) *V. cholerae* and purified twice on the same host. Total gDNA was prepped as above. NEBNext Ultra II DNA Library Preparation Kit for Illuminia (New England Biolabs) was used to prep genomic DNA and was sequenced by paired-end sequencing (2 × 150 bp) on an Illumina HiSeq4000 (University of California, Berkeley QB3 Core Facility). The wild-type phage genome was assembled using SPAdes (Bankevich et al. 2012) with paired-end reads and default settings. This assembly was used as the reference sequence for comparison to escape phage sequence reads with breseq (Deatherage & Barrick 2014) in ‘consensus’ mode and default settings. Protein alignments were analyzed using Praline (Bawono & Heringa 2014). HelA conservation was determined by analyzing HelA from 17 phages isolated between 2001 and 2017 (Angermeyer et al. 2018; McKitterick et al. 2019). Phages that did not have whole genome information were Sanger sequenced from previously prepped phage gDNA (McKitterick et al. 2019) with the primers listed in Table S4. Phages included in the phylogenetic analysis were selected from a BLASTP search of HelA and HelB. Each hit was included if it had over 30% identity to either protein across 90% of the protein. A multiple alignment of helicase amino acid sequences was generated with MUSCLE v3.8.31 (Edgar 2004) using default settings. The alignment file was converted to the PHYLIP format with Clustal X v2.0 (Larkin et al. 2007) and a bootstrapped (n=100) maximum-likelihood phylogenetic tree was solved using PhyML v20120412 (Guindon et al. 2005) with the following settings: -d aa -s BEST --rand_start --n_rand_starts 100 -o tlr -b 100).

#### Quantification and statistical analysis

Statistical tests used for experiments are listed in the Methods section. Data was analyzed using Prism GraphPad. For EOPs, qPCR, lysis kinetics, and nanoluciferase assays, error bars indicate standard deviation of average fold change from three independent biological replicates. Spot plate, agarose gel, and blot images are representative of at least two independent experiments.

#### Data and code availability

The data supporting the study are found in the manuscript, supplementary information, or from the corresponding author upon request.

