## Supplemental figures for "Viral satellites exploit phage proteins to escape degradation of the bacterial host chromosome"

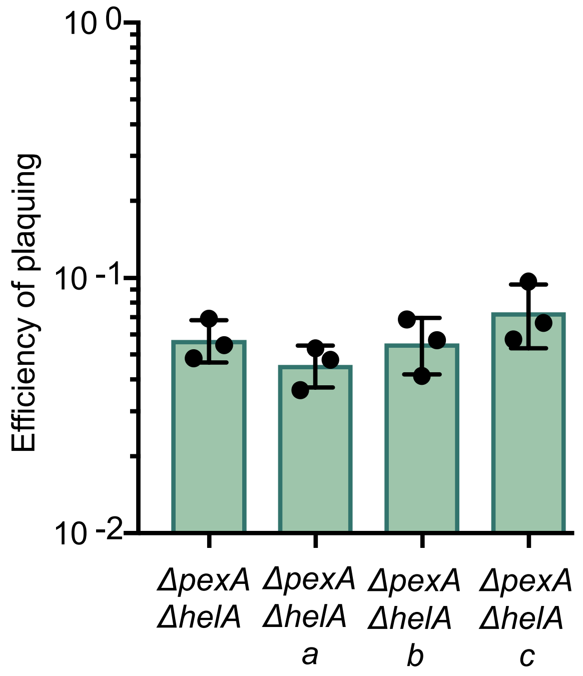


**Figure S1. Related to Figure 1. ICP1[A] *ΔpexA ΔhelA* does not accumulate mutations to escape PLE.** Three sets of plaques (a-c) from ICP1 *ΔpexA ΔhelA* on PLE 1 *V. cholerae* were picked and the efficiency of plaquing on PLE 1 relative to PLE (-) *V. cholerae* was tested.

**A**

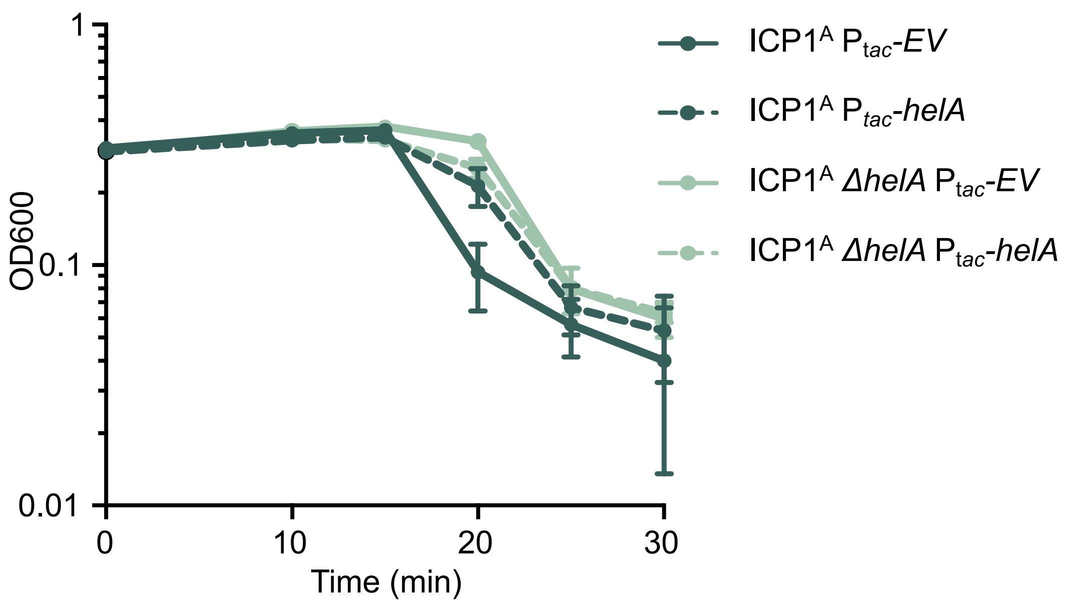


**B**

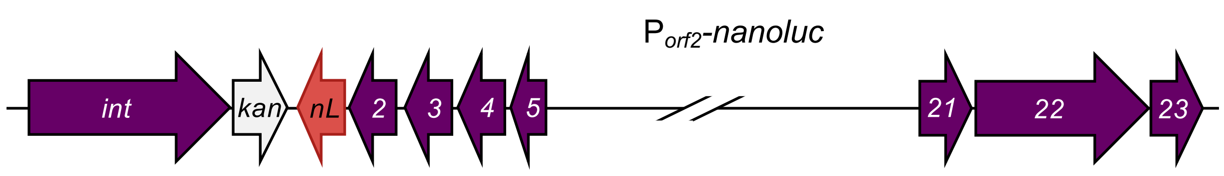


**Figure S2. Related to Figure 2. Infected PLE *1 V. cholerae* demonstrate altered lysis kinetics when PLE 1 does not replicate. A,** $OD_{600}$ of the PLE 1 *V. cholerae* over time after infection with the listed ICP1. **B,** Cartoon of the PLE 1-encoded nanoluciferase reporter with nanoluciferase (nL) encoded downstream of PLE 1 *orf2.* Line break in the PLE genome is shown for simplicity and no other mutations are present.

**A**

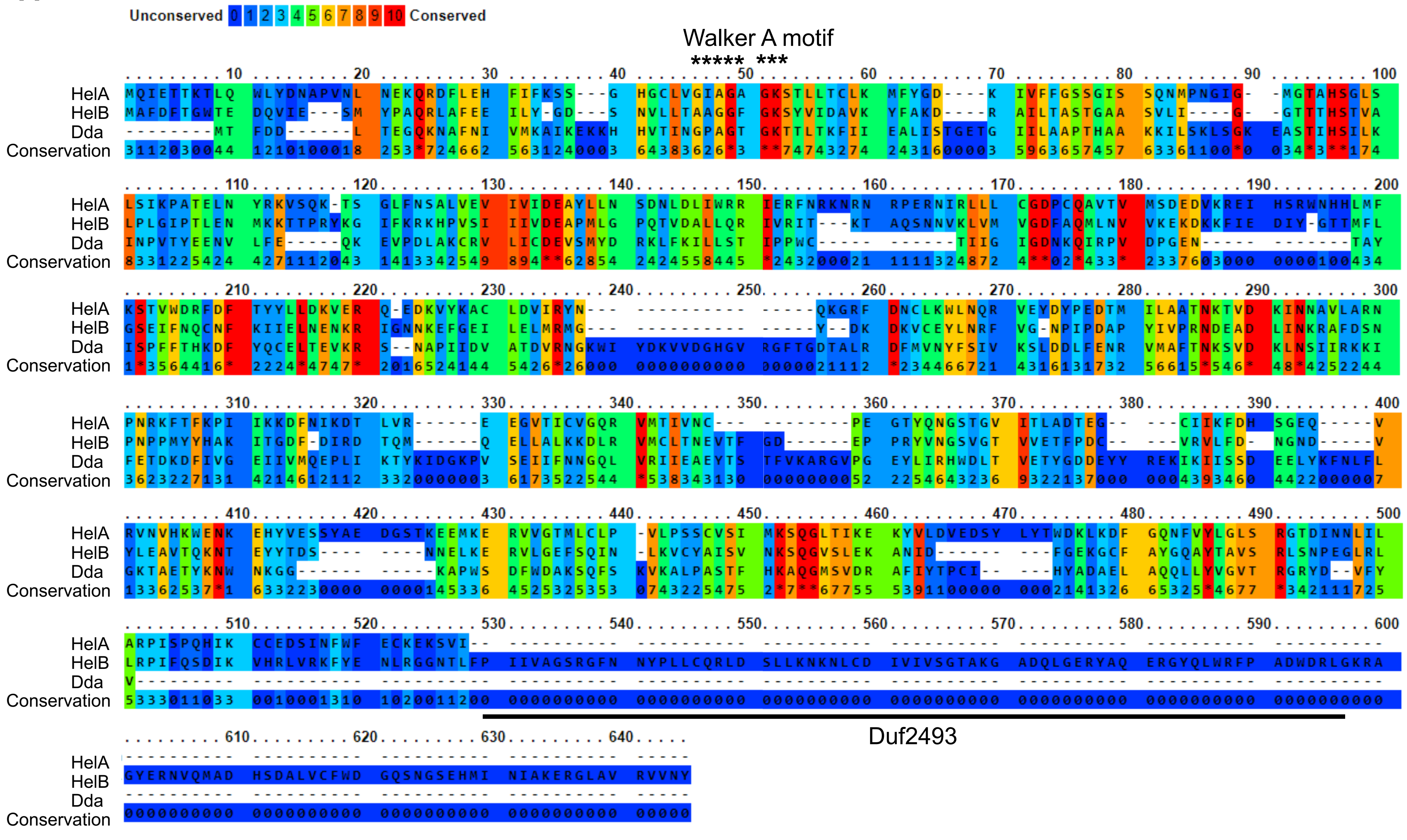


**B**

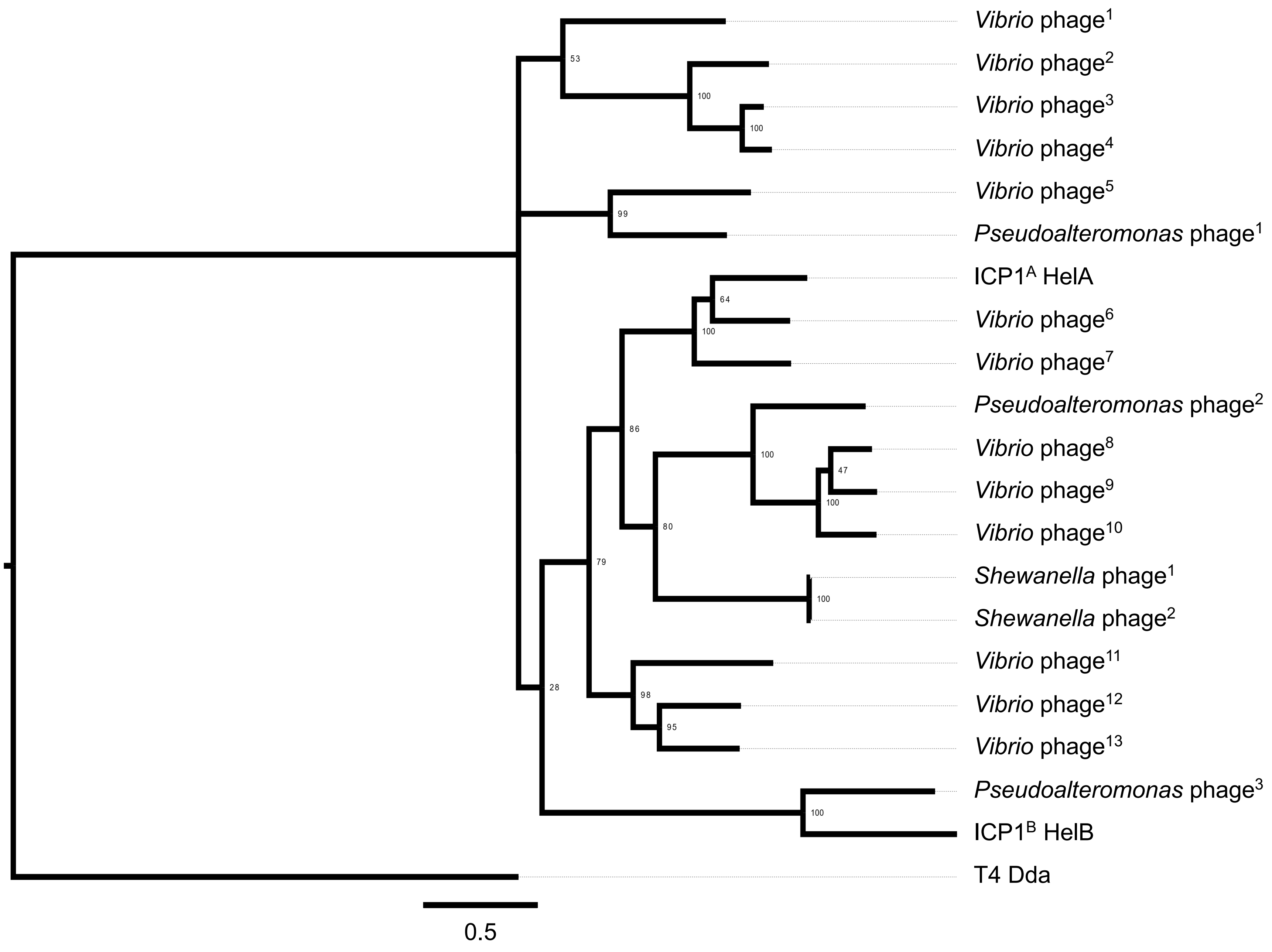


**Figure S3. Related to Figure 4. SF1B-type helicases are found in a variety of marine phages. A,** Praline alignment (Bawono & Heringa 2014) of $ICP1^A$ HelA, $ICP1^B$ HelB, and T4 Dda. The Walker A motif used in ATP hydrolysis is indicated by asterisks, and the Duf2493 in HelB is underlined. **B,** Phylogenetic analysis of SF1B-type helicases, phages used are listed in Table S7.

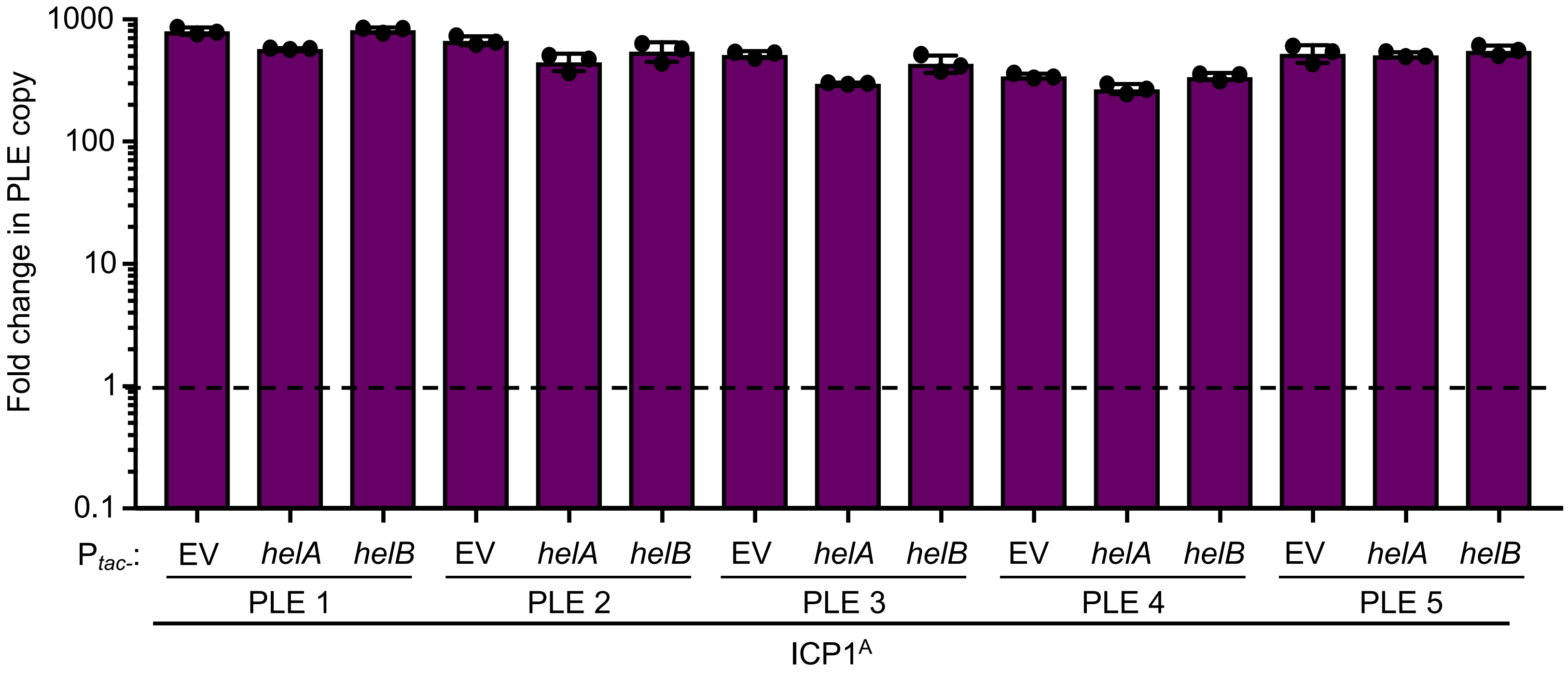


**Figure S4. Related to Figure 6. PLE replication is not altered by ectopic expression of *helA* or *helB*.** Replication of the listed PLE in an isogenic *V. cholerae* background 20 minutes following infection by ICP1$^{A}$. Ectopic vectors were induced 20 minutes prior to infection. Dashed line indicates no change in copy.
