## Supplemental tables for "Viral satellites exploit phage proteins to escape degradation of the bacterial host chromosome"

### Table S1. Related to Figure 1. Escape phage gene products that acquired mutations

| Phage | *gp63* | *gp121* | *gp139* | *gp147* | *gp176* |
|---|---|---|---|---|---|
| **escape Φ1** | | | | Q211E | |
| **escape Φ2** | | | M66I | Δ16 bp | |
| **escape Φ3** | R12C | I321I | | Δ16 bp | A274T |

### Table S2. Strains used in this study

| Bacterial strains | Strain number | Description | Source |
|---|---|---|---|
| **PLE (-)** | KDS6 | *V. cholerae* O1, El Tor biotype; SmR, E7946 | (Levine et al. 1982) |
| **PLE 1** | KDS 36 | E7946 containing PLE 1 integrated into VCR between VCA0329 and VCA0330 | (O'Hara et al. 2017) |
| **midiPLE** | KDS232 | E7946 containing PLE 1 *int*, the PLE 1 origin, and a kanamycin resistance cassette integrated in VCR between VCA0329 and VCA0330; with PLE 1 *repA* cloned into the *lacZ* locus under a $P_{tac}$ promoter and marked with a spectinomycin resistance cassette. | (Barth et al. 2019) |
| **$P_{orf2}$-*nanoluc*** | KDS260 | E7946 containing PLE 1 with nanoluciferase and a kanamycin resistance cassette cloned downstream of *orf2* | This study |
| **miniPLE** | KDS186 | E7946 containing PLE 1 *int* and Kanamycin resistance cassette integrated in VCR between VCA0329 and VCA0330 | (McKitterick & Seed 2018) |
| **$miniPLE_{CD}$** | KDS261 | miniPLE Int S11A | This study |
| **PLE 2** | KDS37 | E7946 containing PLE 2 interrupting VCA0581 | (O'Hara et al. 2017) |
| **PLE 3** | KDS38 | E7946 containing PLE 3 integrated in VCR in between VCA0415 and VCA0416 | (O'Hara et al. 2017) |
| **PLE 4** | KDS39 | E7946 containing PLE 4 integrated in VCR in between VCA0353 and VCA0354 | (O'Hara et al. 2017) |
| **PLE 5** | KDS40 | E7946 containing PLE 3 integrated in VCR in between VCA0407 and VCA0408 | (O'Hara et al. 2017) |

### Table S3. Phages used in this study

| Phage isolates | Strain number | Description | Source |
|---|---|---|---|
| **$ICP1^A$** | KSΦ38 | ICP1_2006_E *ΔCRISPR Δcas2-3* | (McKitterick & Seed 2018) |
| **$ICP1^A$ *ΔpexA*** | ACMΦ142 | ICP1_2006_E *ΔCRISPR Δcas2-3 ΔpexA* | (McKitterick & Seed 2018) |
| **$ICP1^A$ *ΔhelA*** | ACMΦ262 | ICP1_2006_E *ΔCRISPR Δcas2-3 ΔhelA* | This study |
| **$ICP1^A$ *ΔpexA ΔhelA*** | ACMΦ264 | ICP1_2006_E *ΔCRISPR Δcas2-3 ΔpexA ΔhelA* | This study |
| **$ICP1^B$** | ACMΦ259 | ICP1_2017_F_Mathbaria *Δcas2-3;* ICP1 isolate recovered from cholera patient stool collected from ICDDR,B and engineered to be *Δcas2-3* | This study |
| **$ICP1^B$ *ΔhelB*** | ACMΦ262 | ICP1_2017_F_Mathbaria *Δcas2-3 ΔhelB* | This study |
| **$ICP1^B$ B *ΔHD*** | ACMΦ288 | ICP1_2017_F_Mathbaria *Δcas2-3* $\Delta helB^{118\text{-}192}$; ICP1_2017_F_Mathbaria *Δcas2-3* engineered with a deletion between nucleotides 118-192 in *helB* | This study |

### Table S4. Primers used in this study

| Primer | Sequence (5' - 3') | Application |
|---|---|---|
| **Zac14** | AGGGTTTGAGTGCGATTACG | PLE qPCR FWD |
| **Zac15** | TGAGGTTTTACCACCTTTTGC | PLE qPCR RV |
| **Zac68** | CTGAATCGCCCTACCCGTAC | ICP1 qPCR FWD |
| **Zac69** | GTGAACCAACCTTTGTCGCC | ICP1 qPCR RV |
| **Zac10** | ATGCAATGCAGCCATAAACA | miniPLE qPCR FWD |
| **Zac11** | GCGTTTAGTTCGGTGTTGGT | miniPLE qPCR RV |
| **KS364** | CCGCTATCTTTCGAGGTAGC | circularization PCR FWD |
| **KS365** | GCTACTCTCCGTTAAATTCCG | circularization PCR RV |
| **KS587** | ATTCCGGGGATCCGTCGACC | amplify nanoluciferase |
| **KS586** | TGTAGGCTGGAGCTGCTTCG | amplify nanoluciferase |
| **KS324** | GGACGCGAAGCTGGGGATCC | Amplify kanR cassette, DIG Southern probe design |
| **KS325** | ACTCAGGAGAGCGTTCACCG | Amplify kanR cassette, DIG Southern probe design |
| **KS372** | GGTGATTAATTGCTATACAAGTGG | Engineering $P_{orf2}$-*nanoluc* |
| **KS501** | GGATCCCCAGCTTCGCGTCCGCGGTCTTTTTCTTACG TGATTTTAAGC | Engineering $P_{orf2}$-*nanoluc* |
| **KS502** | CGGTGAACGCTCTCCTGAGTCCTATTGAGGCGGTTTA TTTTATGTG | Engineering $P_{orf2}$-*nanoluc* |
| **KS1168** | CGAACGCATTCTGGCGTAAACACATAAAATAAACCGC CTCAATAGG | Engineering $P_{orf2}$-*nanoluc* |
| **KS1169** | GTGAAGACCATTACGTTACCTCCttttatgtgtTTAG GAAACCGTAGACAG | Engineering $P_{orf2}$-*nanoluc* |
| **KS369** | CGTAACTAAATTGGTGGTGTGC | Engineering $P_{orf2}$-*nanoluc* |
| **ACM153** | CTGACATTGATTTCCCTCCG | Engineering $miniPLE_{CD}$ |
| **ACM219** | GGCGTTTACTAGCTACTCGTC | Engineering $miniPLE_{CD}$ |
| **ACM218** | GACGAGTAGCTAGTAAACGCC | Engineering $miniPLE_{CD}$ |
| **ACM26** | CTATGTGAACCAAAAGTTGAGCG | Engineering $miniPLE_{CD}$ |
| **gp58F** | AACGCTGCTTTTCCTTTTGA | ICP1 *DNA pol* detection |
| **gp58R** | CCCAGCATTGAGGACACTTT | ICP1 *DNA pol* detection |
| **ACM295** | TTGCACTTGCAGCAACATGG | ICP1 *helA* detection and sequencing |
| **ACM367** | AAAGCGTTCAATACGACGCC | ICP1 *helA* detection and sequencing |
| **ACM294** | GTTGTGATATGTTCTAGGTGCG | ICP1 *helA* sequencing |
| **ACM531** | GTTTCTACTACTGTACCGAC | ICP1 *helB* detection |
| **ACM583** | TGGTAATCCTATCCCTGATG | ICP1 *helB* detection |

### Table S5. Plasmids used in this study

| Plasmids | Cloning Strain | Description | Source |
|---|---|---|---|
| **$P_{tac}$-*EV*** | ACM707 | pMMB67E plasmid engineered to contain riboswitch A downstream of $P_{tac}$ empty vector control | This study |
| **$P_{tac}$-*helA*** | ACM709 | pMMB67E plasmid engineered to contain riboswitch A downstream of $P_{tac}$, for inducible expression of $ICP1^{A}$ *helA* | This study |
| **$P_{tac}$-*helB*** | ACM711 | pMMB67E plasmid engineered to contain riboswitch A downstream of $P_{tac}$, for inducible expression of $ICP1^{B}$ *helB* | This study |
| **$P_{tac}$-*dda*** | ACM765 | pMMB67E plasmid engineered to contain riboswitch A downstream of $P_{tac}$, for inducible expression of T4 *dda* | This study |

**Table S6. Related to Figure 4. Phage isolates used in HelA sequence analysis**

| Phage | Accession | Source |
|---|---|---|
| **ICP1** | HQ641347 | (Seed et al. 2011) |
| **ICP1_2001_A** | HQ641353 | (Seed et al. 2011) |
| **ICP1_2001_B** | KY883636 | (Naser et al. 2017; Angermeyer et al. 2018) |
| **ICP1_2001_C** | KY883637 | (Naser et al. 2017; Angermeyer et al. 2018) |
| **ICP1_2001_D** | KY065147 | (Naser et al. 2017; Angermeyer et al. 2018) |
| **ICP1_2001_E** | KY883634 | (Naser et al. 2017; Angermeyer et al. 2018) |
| **ICP1_2001_F** | KY883635 | (Naser et al. 2017; Angermeyer et al. 2018) |
| **ICP1_2004_A** | HQ641354 | (Seed et al. 2011) |
| **ICP1_2005_A** | HQ641352 | (Angermeyer et al. 2018) |
| **ICP1_2006_A** | HQ641351 | (Seed et al. 2011) |
| **ICP1_2006_B** | HQ641350 | (Seed et al. 2011) |
| **ICP1_2006_E** | MH310934 | (Angermeyer et al. 2018) |
| **ICP1_2009_A** | KY883638 | (Naser et al. 2017; Angermeyer et al. 2018) |
| **ICP1_2011_A** | MH310933 | (Angermeyer et al. 2018) |
| **ICP1_2011_B** | MH310935 | (Angermeyer et al. 2018) |
| **ICP1_2011_C** | KY883639 | (Naser et al. 2017; Angermeyer et al. 2018) |
| **ICP1_2012_B** | MH310936 | (Naser et al. 2017; Angermeyer et al. 2018) |
| **ICP1_2015_A** | * | (McKitterick et al. 2019) |
| **ICP1_2016_A** | * | (McKitterick et al. 2019) |
| **ICP1_2017_A** | * | (McKitterick et al. 2019) |
| **ICP1_2017_B** | * | (McKitterick et al. 2019) |
| **ICP1_2017_C** | * | (McKitterick et al. 2019) |
| **ICP1_2017_D** | * | (McKitterick et al. 2019) |
| **ICP1_2017_E** | * | (McKitterick et al. 2019) |
| **ICP1_2017_F** | * | (McKitterick et al. 2019) |

*Sequenced genomes not available. *helA* was amplified and sequenced using Sanger Sequencing.

### Table S7. Related to Figure S3. Proteins used in SF1B helicase phylogeny

| Phage isolate | Protein accession | Reference |
|---|---|---|
| **ICP1[A] HelA** | ADX89559 | (Seed et al. 2011) |
| **ICP1[B] HelB** | ADX88195 | (Seed et al. 2011) |
| ***Pseudoalteromonas* phage[1#]** | APC44385 | (Gong et al. 2017) |
| ***Pseudoalteromonas* phage[2]** | ASU03327 | N/A |
| ***Pseudoalteromonas* phage[3]** | YP_009225645 | N/A |
| ***Shewanella* phage[1]** | YP_009104072 | (Luhtanen et al. 2014) |
| ***Shewanella* phage[2]** | YP_009100375 | (Luhtanen et al. 2014) |
| **T4 Dda** | ADJ39734 | (Petrov et al. 2010) |
| ***Vibrio phage[1]*** | AUR92300 | (Kauffman et al. 2018) |
| ***Vibrio phage[2]*** | AUR89265 | (Kauffman et al. 2018) |
| ***Vibrio phage[3]*** | AUR91655 | (Kauffman et al. 2018) |
| ***Vibrio phage[4]*** | YP_007877404 | |
| ***Vibrio phage[5]*** | AUR93419 | (Kauffman et al. 2018) |
| ***Vibrio phage[6]*** | AUR86306 | (Kauffman et al. 2018) |
| ***Vibrio phage[7]*** | YP_007673653 | |
| ***Vibrio phage[8]*** | AUR87502 | (Kauffman et al. 2018) |
| ***Vibrio phage[9]*** | AOQ26819 | |
| ***Vibrio phage[10]*** | AUR8488 | (Kauffman et al. 2018) |
| ***Vibrio phage[11]*** | YP_007676005 | |
| ***Vibrio phage[12]*** | AUR94225 | (Kauffman et al. 2018) |
| ***Vibrio phage[13]*** | BAV80879 | (Ramphul et al. 2017) |

[#]phages are numbered as in Figure S3

### Table S8. Related to Figure 4. Phages probed for helA/B allele geographical distribution

| Phage isolates | Strain number | Collection location | Sample number | Date of Collection[†] | Source |
|---|---|---|---|---|---|
| ***helA* (+); ICP1_2006_E** | KSΦ38 | Dhaka | - | 2006 | (Seed et al. 2011) |
| ***helB* (+); ICP1_2006_D** | KSΦ65 | Dhaka | - | 2006 | (Seed et al. 2011) |
| **ICP1_2017_A** | ACMΦ194 | Dhaka | KDCP7 | 04/02/2017 | (McKitterick et al. 2019) |
| **ICP1_2017_B** | ACMΦ204 | Dhaka | KDCP18 | 04/08/2017 | (McKitterick et al. 2019) |
| **ICP1_2017_C** | ACMΦ197 | Dhaka | KDCP19 | 05/04/2017 | (McKitterick et al. 2019) |
| **ICP1_2017_D** | ACMΦ185 | Dhaka | KDCP24 | 05/16/2017 | (McKitterick et al. 2019) |
| **ICP1_2017_E** | ACMΦ190 | Dhaka | KDCP37 | 05/25/2017 | (McKitterick et al. 2019) |
| **ICP1_2017_F** | ACMΦ199 | Dhaka | KDCP40 | 05/27/2017 | (McKitterick et al. 2019) |
| **ICP1_2017_A_MATHBARIA** | SGHΦ61 | Mathbaria | KMCP3 | 04/20/2017 | This study |
| **ICP1_2017_B_MATHBARIA** | SGHΦ62 | Mathbaria | KMCP8 | 04/24/2017 | This study |
| **ICP1_2017_C_MATHBARIA** | SGHΦ63 | Mathbaria | KMCP9 | 04/24/2017 | This study |
| **ICP1_2017_D_MATHBARIA** | SGHΦ64 | Mathbaria | KMCP10 | 04/26/2017 | This study |
| **ICP1_2017_E_MATHBARIA** | SGHΦ65 | Mathbaria | KMCP12 | 04/26/2017 | This study |
| **ICP1_2017_F_MATHBARIA** | SGHΦ66 | Mathbaria | KMCP13 | 04/27/2017 | This study |

[†]Dates are presented as month/day/year where available
